# Membrane Retrieval after Immediately Releasable Pool (IRP) Exocytosis is produced by Dynamin-Dependent and Dynamin-Independent Mechanisms

**DOI:** 10.1101/2022.08.31.506099

**Authors:** Lucas Bayonés, Mauricio Montenegro, José Moya-Díaz, Samuel Alfonso-Bueno, Luciana I. Gallo, Fernando D. Marengo

## Abstract

The importance of the immediately releasable pool (IRP) of vesicles was proposed to reside in the maintenance of chromaffin cell secretion during the firing of action potentials at basal physiological frequencies. To accomplish this duty, IRP should be replenished as a function of time. We have previously reported that an action potential-like stimulus (APls) triggers the release of ∽50% IRP, followed by a fast dynamin-dependent endocytosis and an associated rapid replenishment process. In this work we investigated the endocytosis and IRP replenishment produced after the exocytosis of variable IRP fractions in mice primary chromaffin cell cultures. Exocytosis and endocytosis were estimated by membrane capacitance measurements obtained in patch-clamped cells. In addition to the dynamin-dependent fast endocytosis activated after the application of APls or 5 ms squared depolarizations, we found that depolarizations lasting 25-50 ms, which release >80% of IRP, are related with a fast dynamin-independent, Ca^2+^- and protein kinase C (PKC)-dependent endocytosis (time constant < 1 s). PKC inhibitors, such as staurosporine, bisindolylmaleimide XI and prolonged treatments with high concentrations of phorbol esters, reduced and decelerated this endocytosis. Additionally, we found that the inhibition of PKC also abolished a slow component of replenishment (time constant ∽8 s) observed after total IRP exocytosis. Therefore, our results suggest that PKC contributes to the coordination of membrane retrieval and vesicle replenishment mechanisms that occur after the complete exocytosis of IRP.

## Introduction

The immediately releasable pool (IRP) is a small group of ready releasable vesicles rapidly released by short depolarizations due to their proximity to voltage dependent Ca^2+^ channels (VDCC) (Alvarez & Marengo 2011; Marengo 2005; Voets *et al*. 1999; Horrigan & Bookman 1994). Due to this characteristic, IRP is believed to be particularly sensitive to sharp gradients of Ca^2+^ developed close to VDCC. Therefore IRP can be important in secretion during action potentials firing at low frequencies where residual Ca^2+^ buildup is supposed to be absent or small (Moya-Diaz *et al*. 2016; Oré & Artalejo 2005).

The replenishment of IRP is essential to maintain the secretion from this pool as a function of time. It was described recently that after complete exocytosis, IRP is replenished by rapid (time constant ∽1 s) and slow (time constant ∽10 s) components (Montenegro *et al*. 2021). The rapid replenishment process can be isolated by the application of a brief depolarization, similar in amplitude and temporal course to a native action potential, which provokes the release of a small fraction of IRP (i.e., ETAP, for exocytosis triggered by action potential) (Moya-Diaz *et al*. 2016; Montenegro *et al*. 2021). This rapid replenishment process accounts for the maintenance of synchronous exocytosis during repetitive stimulation at action potential rates below 0.5 Hz, which is within the range of basal frequencies measured in chromaffin cells (Moya-Diaz *et al*. 2016; Wallace *et al*. 2002; de Diego *et al*. 2008; Marcantoni *et al*. 2010; Holman *et al*. 1994). We showed that ETAP exocytosis is immediately followed by a dynamin-(and F-actin) dependent fast endocytosis, and when any of these two proteins are inhibited or disrupted, fast endocytosis and ETAP replenishment are both decelerated in parallel (Montenegro *et al*. 2021; Moya-Diaz *et al*. 2016). Therefore, we postulated that fast dynamin-dependent endocytosis is essential for ETAP replenishment. However, we do not have information about the membrane retrieval mechanisms operating after the exocytosis of bigger fractions of IRP, which are induced by longer depolarizations than an action potential. Furthermore, we do not have information on the mechanisms associated with the slow replenishment component, activated after the complete IRP release.

It was demonstrated along the years that exocytosis is associated with the activation of different types of endocytosis (Wu *et al*. 2014; Wu *et al*. 2009; Chan & Smith 2001). In secretion terms, it was proposed that the importance of endocytosis is to retrieve the membrane previously fused by exocytosis, to re-internalize vesicle proteins, to restore the structure of the release sites, and to initiate the membrane recycling pathway that ends in the production of new releasable vesicles (Wu *et al*. 2014; Saheki & De Camilli 2012). Several investigators have demonstrated that changes in the stimulation pattern or strength promote transitions between different forms of endocytosis (Artalejo *et al*. 2002; Smith & Neher 1997; Wu *et al*. 2009; Chan & Smith 2001). The goal of this work was to investigate the endocytotic mechanisms operating in the range of stimuli that releases increasing fractions of IRP, and to elucidate the possible relationship of these mechanisms with IRP replenishment processes. Our results demonstrate that, in agreement with our previous data, short stimulus releasing less than 50% of total IRP activates a dynamin-dependent fast endocytosis. More importantly, 25 and 50 ms squared depolarizations, which release approximately 80 and 100% of IRP, are related with a dynamin-independent, but Ca^2+^- and protein kinase C (PKC)-dependent, endocytosis process. Finally, we also found that the slow component of replenishment, observed after total IRP exocytosis (Montenegro *et al*. 2021), is also PKC-dependent. Therefore, our results suggest that PKC coordinates membrane retrieval and vesicle replenishment mechanisms that occur after the complete exocytosis of IRP.

## Materials and Methods

### Mouse Adrenal Chromaffin Cell Preparation

All animal procedures were performed under protocols approved by the Consejo Nacional de Investigaciones Científicas y Técnicas (CONICET, Argentina) and the Facultad de Ciencias Exactas y Naturales (Universidad de Buenos Aires) (CICUAL, experimental protocol N° 065), and are in accordance with the National Institute of Health Guide for the Care and Use of Laboratory Animals (NIH publication 80-23/96), USA. This study was not pre-registered. One hundred ninety two 129/sv wild type mice (13-21 days old, both sexes) were obtained from the Bioterio Central (Facultad de Ciencias Exactas y Naturales, Universidad de Buenos Aires, Argentina). No other inclusion or exclusion criteria were pre-determined for the mice. Animals were housed (6-8 per cage) with ad libitum access to chow and water, under temperature-controlled conditions on a 12 hours light/dark cycle. Animals were sacrificed with intraperitoneal injection of an overdose (500 mg/kg) of tribromoethanol (avertin) to minimize suffering. The avertin solution (20 mg/ml) was prepared by mechanical agitation (4 hours) at room temperature, and kept at 4 °C until use. This anesthesia provokes the total lack of animal reaction to stimuli in less than 5 min after application (0.5 ml/animal), without any sign of suffering or paradoxical side effects. In general the animals died because of the anesthesia, but to avoid any possible suffering we always performed cervical dislocation immediately after the end of surgery. Adrenal glands from two female/male mice were used in each culture (Alvarez *et al*. 2008). Chromaffin cells were isolated and cultured following the procedures described by Perez Bay et al. (Perez Bay *et al*. 2012). Briefly, mechanically isolated adrenal medullas were digested for 25 min in Hanks solution with papain (0.5– 1 mg/ml) at 37°C, and mechanically disrupted by pipetting through a yellow tip in Dulbecco’s modified Eagle’s medium low glucose, supplemented with 5% fetal calf serum, 5 μl/ml penicillin/streptomycin, 1.3 μl/ml gentamycin, 1 mg/ml bovine serum albumin, and 10 μM citosine-1-β-D-arabinofuranoside. The cell suspension was filtered through 200 and 50 μm pore meshes, and cultured on poly-L-lysine pretreated coverslips at 37°C, 95% O_2_ – 5% CO_2_. The cells were used for experiments between 24-48 hours after culture during daytime (9-20 hs). Transient transfections with the plasmid dynamin I K44A-EGFP (a dynamin-I GTPase-defective dominant-negative mutant) were performed using a Nucleofector 4D device (Lonza, Switzerland), according to the manufacturer’s instructions.

### Whole Cell Patch-Clamp and Membrane Capacitance Measurements

The patch-clamp setup comprised an inverted microscope (Olympus IX71, Olympus, Japan), a patch-clamp amplifier (EPC10 double patch clamp amplifier, Heka Elektronik, Lambrecht, Germany) and a personal computer. The application of the stimulation protocols and the acquisition of data were controlled by the Patchmaster® software (Heka, Lambrecht, Germany, RRID:SCR_000034). Chromaffin cells were washed in a standard extracellular solution composed of (mM) 120 NaCl, 20 Hepes, 4 MgCl_2_, 5 CaCl_2_, 5 mg/ml glucose and 1 μM tetrodotoxin (pH 7.3). The patch-clamp electrodes (3–5 MΩ) were filled with an internal solution containing (mM) 95 Cs D-glutamate, 23 HEPES, 30 CsCl, 8 NaCl, 1 MgCl_2_, 2 Mg-ATP, 0.3 Li-GTP, and 0.5 Cs-EGTA (pH 7.2). These solutions were designed to selectively measure voltage dependent Ca^2+^ currents (I_Ca2+_) and to maximize the exocytosis of vesicles tightly coupled to voltage-dependent Ca^2+^ channels, and were used frequently before by us to study the exocytosis of IRP (Moya-Diaz *et al*. 2016; Alvarez *et al*. 2008). In some experiments the external or internal solutions were changed as indicated in the text. The micropipettes were coated with dental wax to minimize their stray capacitance and to achieve a better C-fast compensation. The holding potential of −80 mV was not corrected for junction potentials. We considered that the recorded cells were “leaky,” and discarded, when the leak current measured at the holding potential was bigger than −10 pA. We also discarded cells with series resistance smaller than 12 MΩ. The cell capacitance (Cm) was measured continuously during the experiments by application of the sine+dc method (Gillis 1995) implemented through the lock-in extension of Patchmaster, using a sinusoidal voltage (800 Hz, 60 mV peak to peak) added to the holding potential. The application of the sinusoidal voltage was suspended from 10 ms before to 10 ms after the depolarizations applied to stimulate the cells. We always worked with the same experimental chamber and with the same volume of external solution (0.5 ml), which was maintained stable during the entire experiment.

### Experimental Design, Data Analysis and Statistics

The exocytosis of different fractions of IRP was induced by single 5, 10, 25 and 50 ms square depolarizations (SQP5ms, SQP10ms, SQP25ms and SQP50ms), from the holding potential of −80 to −10 mV. We know from previous work that, in our standard conditions, a 50 ms depolarization completely releases IRP in our chromaffin cell preparation. We also stimulated the cells with an action potential-like stimulus (APls) composed by a 2.5 ms ascendant voltage ramp from −80 to +50 mV, followed by a 2.5 ms descendant ramp that returns to holding potential. In some cells, immediately after the application of a depolarization pulse, we noted the presence of a brief transient capacitance change, probably associated to sodium channels gating (Horrigan & Bookman 1994). This transient became negligible in less than 50 ms after the end of depolarization (Moya-Diaz *et al*. 2016). A capacitive artifact with similar kinetics was reported before in bovine and embryonic mouse chromaffin cells (Chow *et al*. 1996; Gong *et al*. 2005). Therefore, to avoid any influence of this fast capacitance transient, the synchronous increase in Cm (ΔCm_exo_), associated with exocytosis, was defined as the difference between the averaged Cm measured in a 50 ms window starting 60 ms after the end of the stimulus minus the average pre-stimulus Cm also measured in a 50 ms window (Supplementary Figure 1). In a similar way, the following decreased in capacitance (ΔCm_endo_), associated with endocytosis, was calculated as the difference between the averaged Cm measured in a 50 ms window starting 60 ms after the end of the stimulus minus the averaged Cm measured 5 s after the stimulus, also in a 50 ms window (Supplementary Figure 1). To study the replenishment of IRP we applied a protocol composed of pairs of SQP50ms, where the pulses of each pair are separated by time intervals (Δt) of 0.2, 0.4, 1, 2, 3, 5, 7, 10, 20 and 40 s respectively (Montenegro *et al*. 2021). All these pairs of pulses were repeated in each single cell and applied at random, using a table of random numbers. We expressed the degree of replenishment as the ratio between exocytosis evoked by the second stimulus (ΔCm_2_) and exocytosis evoked by the first stimulus (ΔCm_1_) (Supplementary Figure 14). The data were filtered at 3 kHz. The experiments were carried out at room temperature (22–24°C).

Data are expressed as mean values±standard error of measurements obtained in different cells (one measurement per cell, and one cell per petry dish). For every experimental condition, the data were always obtained from several cell cultures. The number of cell cultures used in every experimental condition and for each measured variable is represented in Supplementary Table 1. No blinding and no sample calculation were performed in this study. No test for outliers was performed. Statistical analysis was performed with the statistical package of Sigma Plot 11.0 (Systat Software, RRID:SCR_003210). We used the Student’s “t” test for comparisons between two groups of independent data samples, and one way ANOVA for multiple independent data samples followed by Bonferroni test for comparisons between groups. We performed the Shapiro-Wilk test to evaluate the normality of samples. To fit the endocytosis records or IRP, or ETAP, replenishment curves, we used the nonlinear curve-fitting option in Origin Pro 8 (Microcal Software, RRID:SCR_002815). The comparison between fittings with different number of exponential components was performed with a Fisher test, which considers the residual sum of square errors and the degree of freedom of each model (Origin statistical package).

## Materials

Bovine serum albumin (CAT No. 05470-25G), poly-L-lysine (CAT No. P8920-100ML), cytosine-1-beta-D-arabinofuranoside (CAT No. C6645-25MG), papain (CAT No. P4762-100MG), Mg ATP (CAT No. A9187-100MG), Li GTP (CAT No. G5884-25MG), EGTA (CAT No. E4378-10G), GTPγS (G8634-1MG), bisindolimaleimide XI (BIS XI) (B4056-1MG) and phorbol 12-myristate 13-acetate (PMA) (P8139-1MG) were obtained from Sigma (St Louis, MO, USA); Dulbecco’s modified Eagle’s medium (CAT No. 10567-014), gentamycin (CAT No. 15750078) and penicillin/streptomycin (CAT No. 15140122) from Gibco (Carlsbad, CA, USA); fetal calf serum from Natocor (Córdoba, Argentina, CAT No. SFB500ml); tetrodotoxin citrate (CAT No. T-550) and staurosporine (S-350) from Alomone Labs (Jerusalem, Israel); and the monoclonal anti-dynamin antibody (BD Biosciences, San Jose, CA, EEUU, CAT No. 610246, RRID:AB_397641).

## Results

### The speed of the endocytosis induced after IRP exocytosis changes with Ca^2+^ entry in a biphasic manner

To induce IRP exocytosis, we reproduced the standard conditions used in our previous publications (Alvarez *et al*. 2008; Moya-Diaz *et al*. 2016). To maximize the exocytosis of vesicles tightly coupled to voltage-dependent Ca^2+^ channels, external and internal solutions contained 5 mM Ca^2+^ and 0.5 mM EGTA respectively (Moya-Diaz *et al*. 2016) (see Materials and Methods). Increasing fractions of IRP were released by the application of an action potential-like stimulus (APls), and squared depolarization pulses of 5 (SQP5ms), 10 (SQP10ms), 25 (SQP25ms), and 50 ms (SQP50ms), respectively. We have previously demonstrated that SQP50ms completely releases IRP in our chromaffin cell preparation in these experimental conditions (Alvarez *et al*. 2013). Figure 1A (ii) and (iii) illustrate typical recordings of Ca^2+^ current (I_Ca2+_) and cell membrane capacitance (ΔCm), respectively, obtained in response to application of APls, SQP5ms and SQP50ms (represented in Figure 1A (i)). The amplitude (I_Ca2+_) and integral (∫I_Ca2+_) of Ca^2+^ currents are summarized in the Supplementary Figure 2A and B, respectively. Although APls provoked the biggest I_Ca2+_ for all these stimuli, ∫I_Ca2+_ increased progressively from APls to squared depolarizations with increasing durations, reaching statistical difference for SQP25ms and SQP50ms. Like ∫I_Ca2+_, the amplitude of exocytosis (ΔCm_exo_) increased in the same direction, from APls to SQP50ms (Figure 1B). In fact, ΔCm_exo_ followed a progressive and saturating increase with ∫I_Ca2+_ (Supplementary Figure 2C, we averaged both parameters for every stimuli), as was previously demonstrated for IRP exocytosis (Alvarez *et al*. 2013; Horrigan & Bookman 1994; Marengo 2005). On the other hand, the amplitude of endocytosis (ΔCm_endo_) showed no changes with the duration of stimuli (Figure 1C) or ∫I_Ca2+_ (Supplementary Figure 2D), and consequently the efficiency of endocytosis, represented by endo/exo (i.e., ratio between ΔCm_endo_ and ΔCm_exo_), decreased (Figure 1D and Supplementary Figure 2E), indicating that in our experimental conditions fast endocytosis cannot retrieve the totality of membrane fused when IRP is completely released. Finally, the time constant of fast endocytosis (τ_endo_) showed an interesting biphasic behavior: increasing its value (i.e., decelerating the endocytosis) between APls and SQP5ms, and decreasing its value (i.e., accelerating the endocytosis) at SQP50ms (Figure 1E). This biphasic behavior is also observed when τ_endo_ is represented versus the increase in ∫I_Ca2+_ (Supplementary Figure 2F). The effect of accumulated ∫I_Ca2+_ on endocytosis was reanalyzed by application of one, two or three consecutive SQP25ms at 13 Hz (Supplementary Figure 3). This protocol provoked approximately proportional increases of ∫I_Ca2+_ with the number of SQP25ms applied (Supplementary Figure 3C). As expected from observation of Figure 1E and the Supplementary Figure 2F, τ_endo_ was inversely correlated with the increase in stimulation intensity and in accumulated ∫I_Ca2+_ (Supplementary Figure 4E and F).

**Figure 1:**
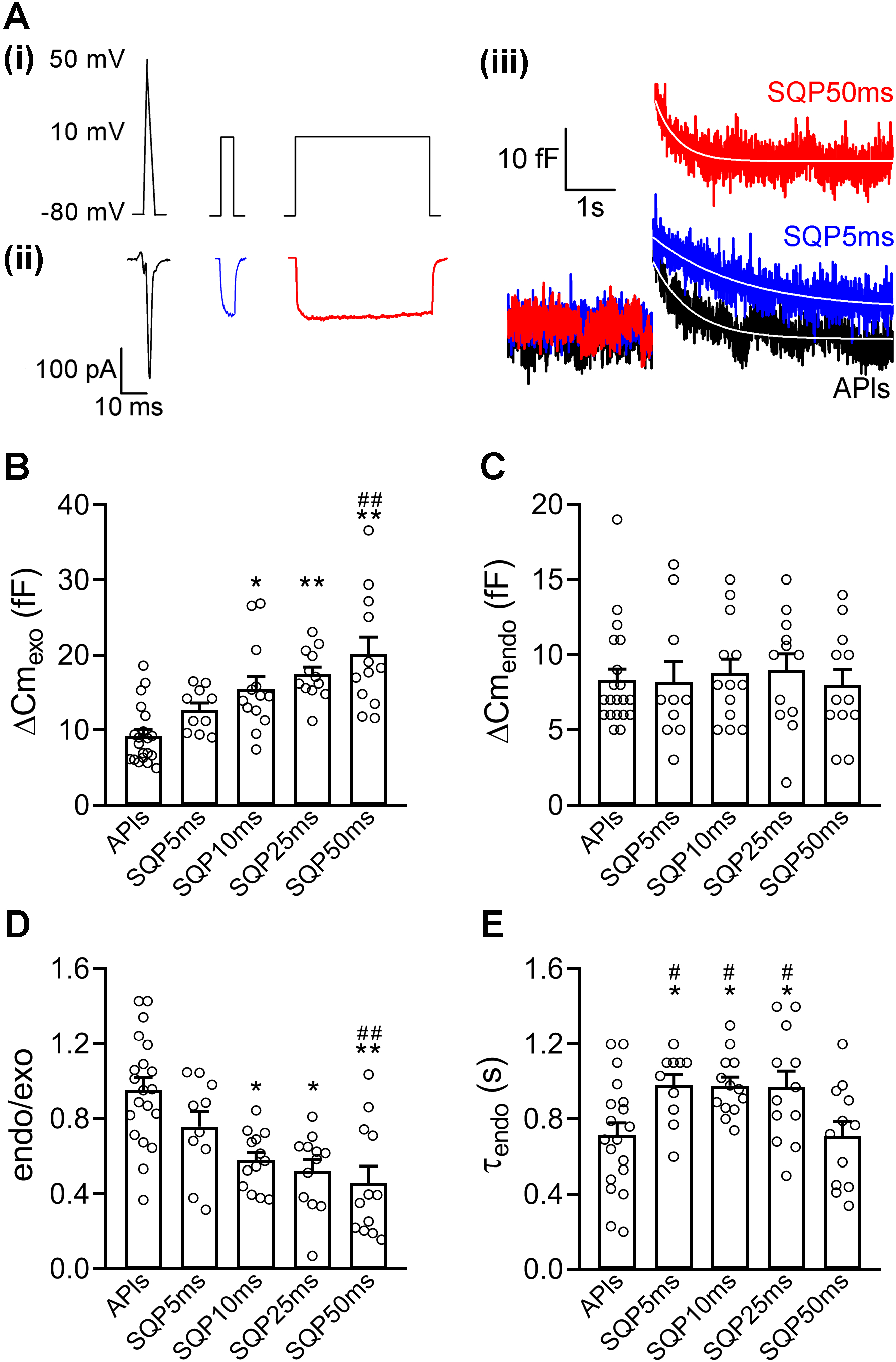
Endocytosis induced after increasing fractions of IRP exocytosis: (A) Schematic representation of APls, SQP5ms and SQP50ms (i), and representative examples of Ca^2+^ currents (ii) and Cm recordings (iii) obtained in response to the application of these stimuli. The white lines in (iii) represent single exponential fittings 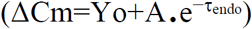 applied to the decay in Cm, where τ_endo_ was 0.74 s (R^2^>0.7557), 0.98 s (R^2^>0.7290) and 0.60 s (R^2^>0.7054) for APls, SQP5ms and SQP50ms, respectively. (B), (C), (D) and (E) The plots represent average values, standard errors and individual measurements (one measurement/cell) of exocytosis (ΔCm_exo_), endocytosis (ΔCm_endo_), ratio between endocytosis and exocytosis (endo/exo), and time constant of endocytosis (τ_endo_) obtained by application of APls, SQP5ms, SQP10ms, SQP25ms and SQP50ms, respectively. The data were analyzed by one-way ANOVA followed by Bonferroni comparisons (* or # p<0.05, ** or ## p<0.01). For figures B and D, * symbols represent comparison of every condition versus APls, and # symbols represent comparison of every condition versus SQP5ms. For figure E, * symbols represent comparison of every condition versus APls, and # symbols represent comparison of every condition versus SQP50ms. The sample sizes (number of individual cells) and the number of cell culture preparations from where these cells were obtained are summarized in Supplementary Table 1 (first line, Control).

To further analyze the effect of Ca^2+^ entry on τ_endo_, we modified the Ca^2+^ concentration of the external solution ([Ca^2+^]_o_). Separate experiments were performed at 1, 5 and 10 mM [Ca^2+^]_o_, while stimulating the cells with SQP50ms. We compared all the variables measured at 1 and 10 mM [Ca^2+^]_o_ versus 5 mM [Ca^2+^]_o_ (our standard condition throughout the study). At both extremes of [Ca^2+^]_o_ tested, I_Ca2+_ and ∫I_Ca2+_ were significantly modified with respect to 5 mM [Ca^2+^]_o_ (Supplementary Figure 5), and τ_endo_ was also changed correspondingly (Figure 2A and E). In other words, τ_endo_ followed an inverse relationship (i.e, the endocytosis is accelerated) with ∫I_Ca2+_ (Figure 2F).

**Figure 2:**
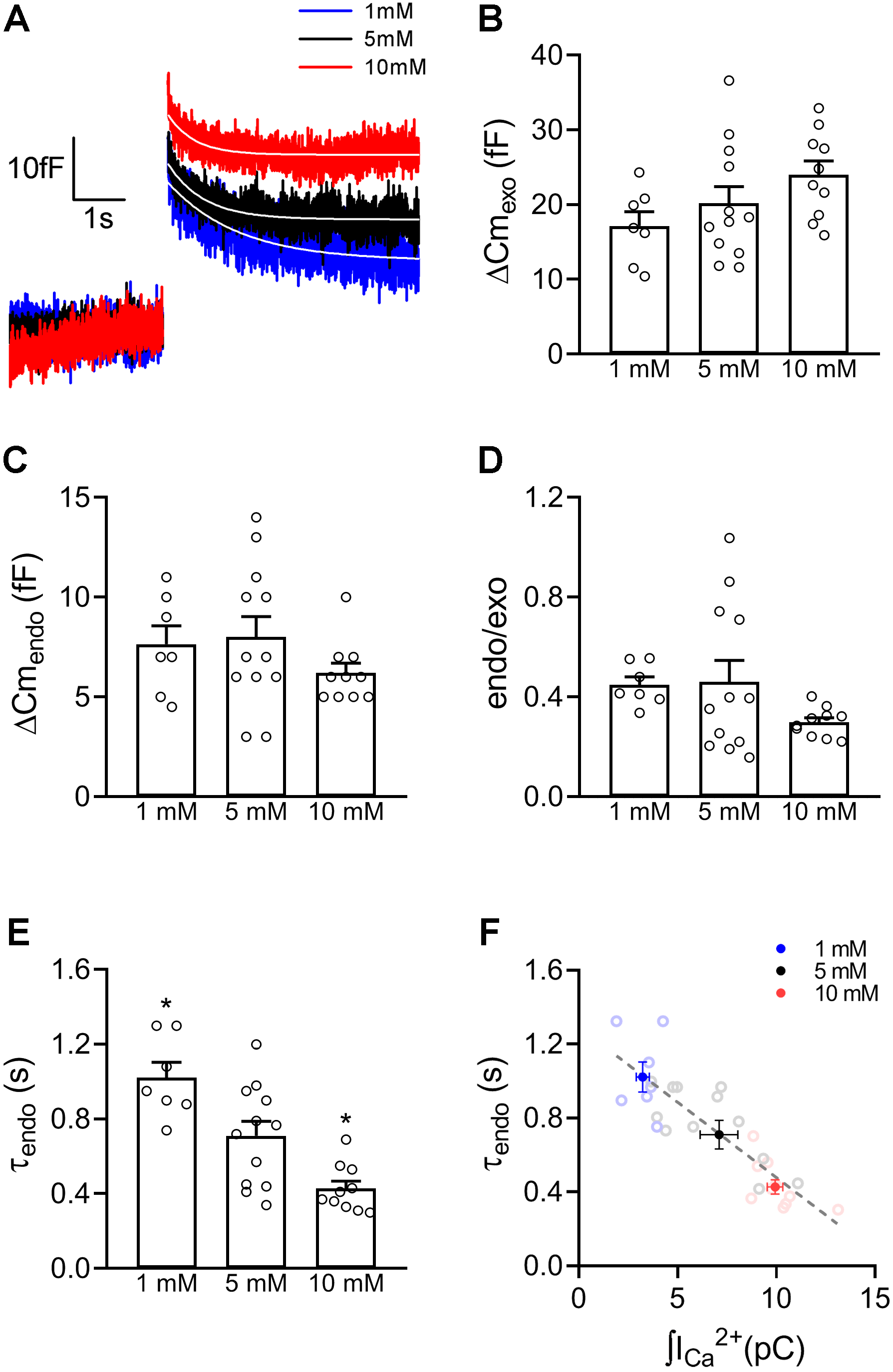
Endocytosis induced after IRP exocytosis with different levels of Ca^2+^ entry: (A) Representative examples of Cm recordings obtained in response to the application of SQP50ms in the presence of 1, 5 and 10 mM [Ca^2+^]_o_ in three independent cells. The white lines represent single exponential fits 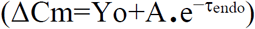 applied to the decay in Cm, where τ_endo_ was 1.38 s (R^2^>0.8284), 0.69 s (R^2^>0.7637) and 0.52 s (R^2^>0.6183) for 1, 5 and 10 mM Ca^2+^, respectively. (B), (C), (D) and (E), The plots represent the average values, standard errors and individual measurements (one measurement/cell) of exocytosis (ΔCm_exo_), endocytosis (ΔCm_endo_), ratio between endocytosis and exocytosis (endo/exo), and time constant of endocytosis (τ_endo_) obtained by application of SQP50ms with 1, 5 and 10 mM [Ca^2+^]_o_ in the external solution. The data were analyzed by one-way ANOVA followed by Bonferroni comparisons. *p<0.05 represents the comparison of every condition versus 5 mM Ca^2+^. (F) The plot represents the average values (closed circles in blue, black and red) and standard errors of τ_endo_ in y-axis versus ∫I_Ca2+_ in x-axis, obtained by application of SQP50ms on cells at 1, 5 and 10 mM [Ca^2+^]_o_, respectively. The dotted line represents a linear fit (τ_endo_=a+b.∫I_Ca2+_) obtained from individual data points (open circles) acquired in the three experimental conditions, where a=1.27, b= −0.08, and R^2^>0.7403. The sample sizes (number of individual cells) and the number of cell culture preparations from where these cells were obtained are summarized in Supplementary Table 1.

This first section of results shows that the rate of endocytosis induced after graded IRP exocytosis is dependent on Ca^2+^ entry. It is important to note that in the two last experimental protocols ΔCm_exo_ did not change between the different experimental conditions (Figure 2B and Supplementary Figure 4), probably because IRP exocytosis is close to saturation (Alvarez *et al*. 2013), indicating that the observed changes in τ_endo_ are not dependent on the magnitude of exocytosis.

The biphasic behavior of τ_endo_ with ∫I_Ca2+_ can be interpreted in different ways. First, it might be a unique endocytic mechanism that switches its Ca^2+^ dependency with [Ca^2+^]. We know from previous work of our laboratory that the fast endocytosis after APls is strongly dependent on dynamin (Moya-Diaz *et al*. 2016). This GTPase is activated/inactivated by de-phosphorylation/phosphorylation processes (Ferguson & De Camilli 2012) that can be, in turn, regulated by Ca^2+^ (Liu *et al*. 1994; Lai *et al*. 1999). A second possibility is that two different processes might contribute to membrane retrieval in different ranges of Ca^2+^ entry. To distinguish between these two possibilities, in the next section we analyze the participation of dynamin in the whole range of stimuli used in this work.

### Membrane retrieval after the complete IRP release is produced through a dynamin-independent endocytosis

To evaluate the dynamin dependence of endocytosis we applied various experimental strategies (see Figures 3, 4, 5 and 6, and Supplementary Figures 6, 7, 8 and 9) on chromaffin cells stimulated by the same multiple stimuli protocol described above (i.e., APls, SQP5ms, SQP10ms, SQP25ms and SQP50ms). In independent experimental series, we applied the GTP non-hydrolyzable analogue GTPγS (Xu *et al*. 2008); intracellular dialysis with a monoclonal antibody against dynamin (Anti-Dyn), that was proved by us and other authors to inhibit the actions of dynamin (Gonzalez-Jamett *et al*. 2010; Montenegro *et al*. 2021; Moya-Diaz *et al*. 2016; Moya-Diaz *et al*. 2020); intracellular dialysis with GST-Dyn829-842 peptide (GST-Dyn), containing the recognition motif for SH3 domain-containing proteins of the dynamin-1 proline-rich domain, which consequently disrupts dynamin-dependent endocytosis (Shupliakov *et al*. 1997; Fulop *et al*. 2008; Moya-Diaz *et al*. 2016); and finally, the expression (by transient transfection) of dynamin-I dominant-negative mutant dynamin-I K44A (DynI K44A), that cannot bind nor hydrolyze GTP (McMahon & Boucrot 2011). Since Anti-Dyn was dialyzed during 5 min before starting the measurements, we performed three different controls: dialysis with normal solution (without Anti-Dyn) during 1 min (Control), dialysis with normal solution during 5 min (DIAL5min), and dialysis with a solution containing a heat-inactivated Anti-Dyn (HI Anti-Dyn) during 5 min. GST-Dyn was also dialyzed during 5 min, so the controls performed were a dialysis with normal solution (Control), and a dialysis with internal solution containing GST alone (GST), both during 5 min before starting the measurements. Finally, the controls for the experiments with Dyn-K44A were mock non-transfected cells (they received the transfection treatment without the plasmid) (Control) and cells transfected with EGFP alone (EGFP). All these treatments produced similar effects on all the measured parameters. None of them affected I_Ca2+_, ∫I_Ca2+_, or ΔCm_exo_ (see the summary of the data in Supplementary Figures 6, 7, 8 and 9, and for examples of ΔCm recordings see Figures 3A, 4A, 5A and 6A, respectively). However, they consistently reduced the amplitude of endocytosis (ΔCm_endo_) and the endo/exo ratio only when APls and SQP5ms were applied (Figures 3B-C, 4B-C, 5B-C, 6B-C), but these treatments did not affect the same parameters for longer stimuli. Additionally, all these treatments increased τ_endo_ (i.e., decelerated endocytosis) for APls and SQP5ms, but there were no modifications to this parameter for the rest of stimuli applied (Figures 3D, 4D, 5D and 6D). The effects observed on the endocytosis associated with these short stimuli are consistent with our previous results (Moya-Diaz *et al*. 2016). However, the most interesting and novel information obtained from these experiments is that none of these treatments affected the amplitude nor the rate of endocytosis observed when SQP10ms, SQP25ms or SQP50ms were applied, indicating the presence of a dynamin-independent endocytosis within this range of stimuli. The Figure 7 represents the re-plotting of τ_endo_ versus ∫I_Ca2+_ for the four experimental series described above. It is interesting to note that while all control conditions show the typical biphasic behavior mentioned before (see Supplementary Figure 2), τ_endo_ consistently decreased with ∫I_Ca2+_ in the experiments where dynamin activity was inhibited by the different treatments. These results suggest that this putative dynamin-independent endocytosis is accelerated with Ca^2+^ entry, in coherence with the results represented in Figure 2F.

**Figure 3:**
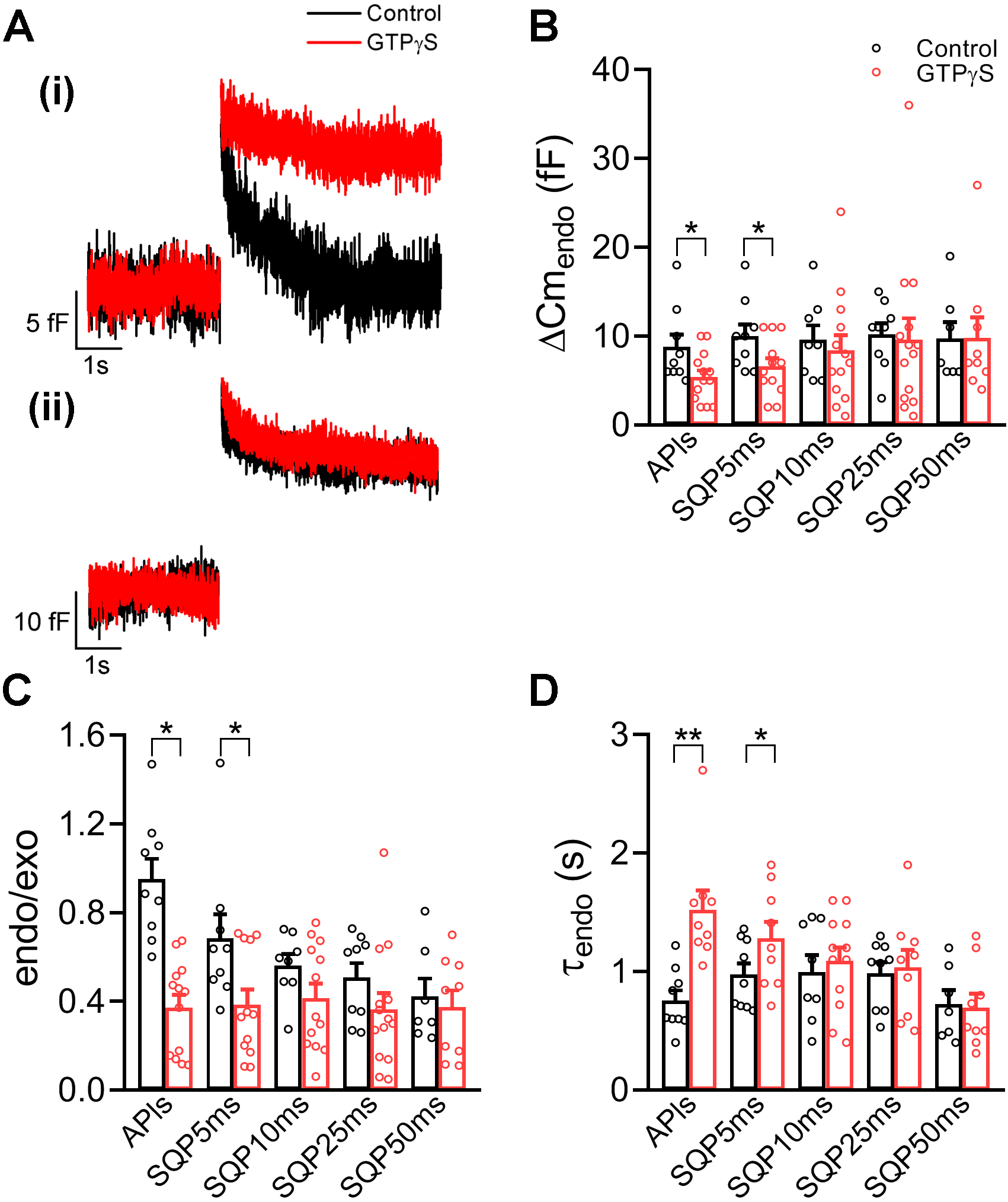
Intracellular application of GTPγS inhibits the endocytosis induced by APls and SQP5ms but not by longer stimuli: (A), Representative examples of Cm recordings obtained in response to the application of APls (i) and SQP50ms (ii) in two independent cells dialyzed with standard internal solution (Control, black) or in presence of 0.3 mM GTPγS (red). (B), (C) and (D), The plots represent average values, standard errors and individual measurements (one measurement/cell) of ΔCm_endo_, endo/exo ratio and τ_endo_ obtained by application of APls, SQP5ms, SQP10ms, SQP25ms and SQP50ms in Control (black) and GTPγS (red) conditions. The data were analyzed by Student’s ‘t’ test. *p<0.05, **p<0.01. The sample sizes (number of individual cells) and the number of cell culture preparations from where these cells were obtained are summarized in Supplementary Table 1.

**Figure 4:**
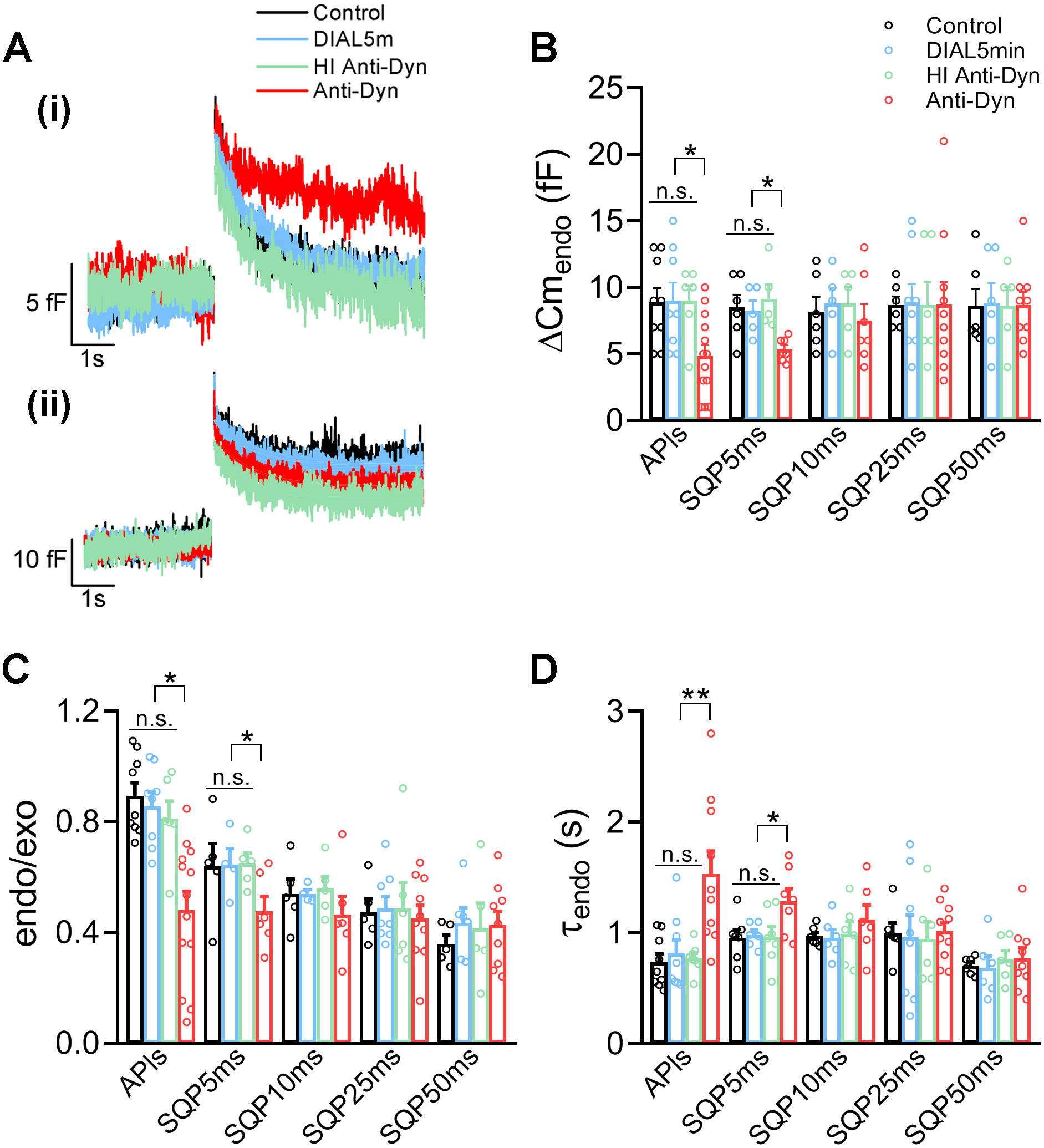
Intracellular application of a monoclonal anti-dynamin antibody (Anti-Dyn) inhibits the endocytosis induced by APls and SQP5ms but not by longer stimuli: (A), Representative examples of Cm recordings obtained in response to the application of APls (i) and SQP50ms (ii) in four independent cells dialyzed with standard internal solution for 1 min (Control, black), for 5 min (DIAL5min, blue), with Anti-Dyn for 5 min (Anti-Dyn, red) or with heat inactivated Anti-Dyn for 5 min (HI Anti-Dyn, green). (B), (C) and (D), The plots represent average values, standard errors and individual measurements (one measurement/cell) of ΔCm_endo_, endo/exo ratio and τ_endo_ obtained by application of APls, SQP5ms, SQP10ms, SQP25ms and SQP50ms in Control (black), DIAL5min (blue), Anti-Dyn (red) and HI Anti-Dyn (green) conditions. The data were analyzed by one-way ANOVA followed by Bonferroni comparisons. *p<0.05, **p<0.01. The sample sizes (number of individual cells) and the number of culture preparations from where these cells were obtained are summarized in Supplementary Table 1.

**Figure 5:**
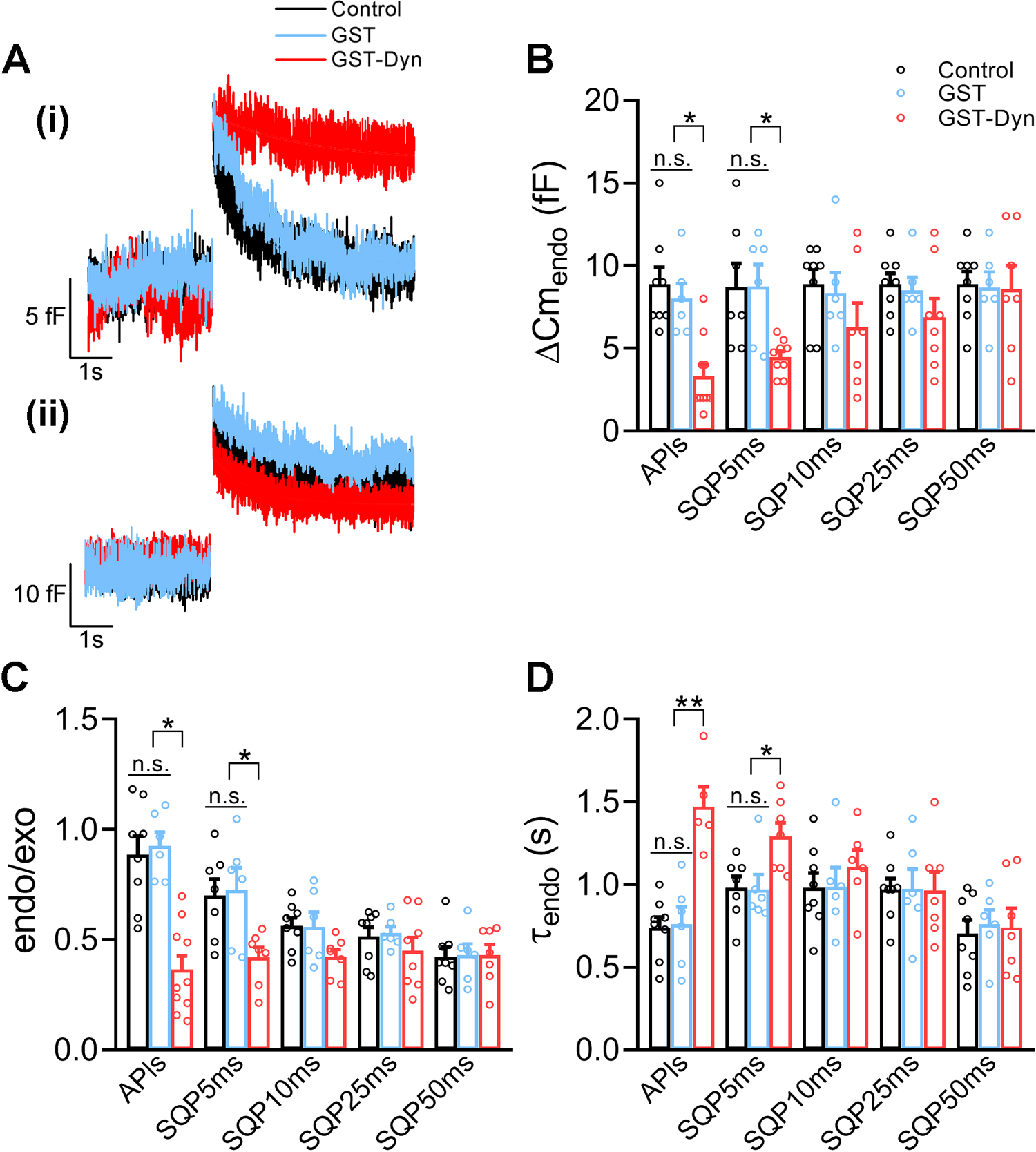
Intracellular application of the peptide GST-Dyn829-842 (GST-Dyn) inhibits the endocytosis induced by APls and SQP5ms but not by longer stimuli: (A), Representative examples of Cm recordings obtained in response to the application of APls (i) and SQP50ms (ii) in three independent cells dialyzed with standard internal solution for 5 min (Control, black), with GST-Dyn for 5 min (GST-Dyn, red) or with GST for 5 min (GST, blue). (B), (C) and (D), The plots represent average values, standard errors and individual measurements (one measurement/cell) of ΔCm_endo_, endo/exo ratio and τ_endo_ obtained by application of APls, SQP5ms, SQP10ms, SQP25ms and SQP50ms in Control (black), GST-Dyn (red) and GST (blue). The data were analyzed by one-way ANOVA followed by Bonferroni comparisons. *p<0.05, **p<0.01. The sample sizes (number of individual cells) and the number of culture preparations from where these cells were obtained are summarized in Supplementary Table 1.

**Figure 6:**
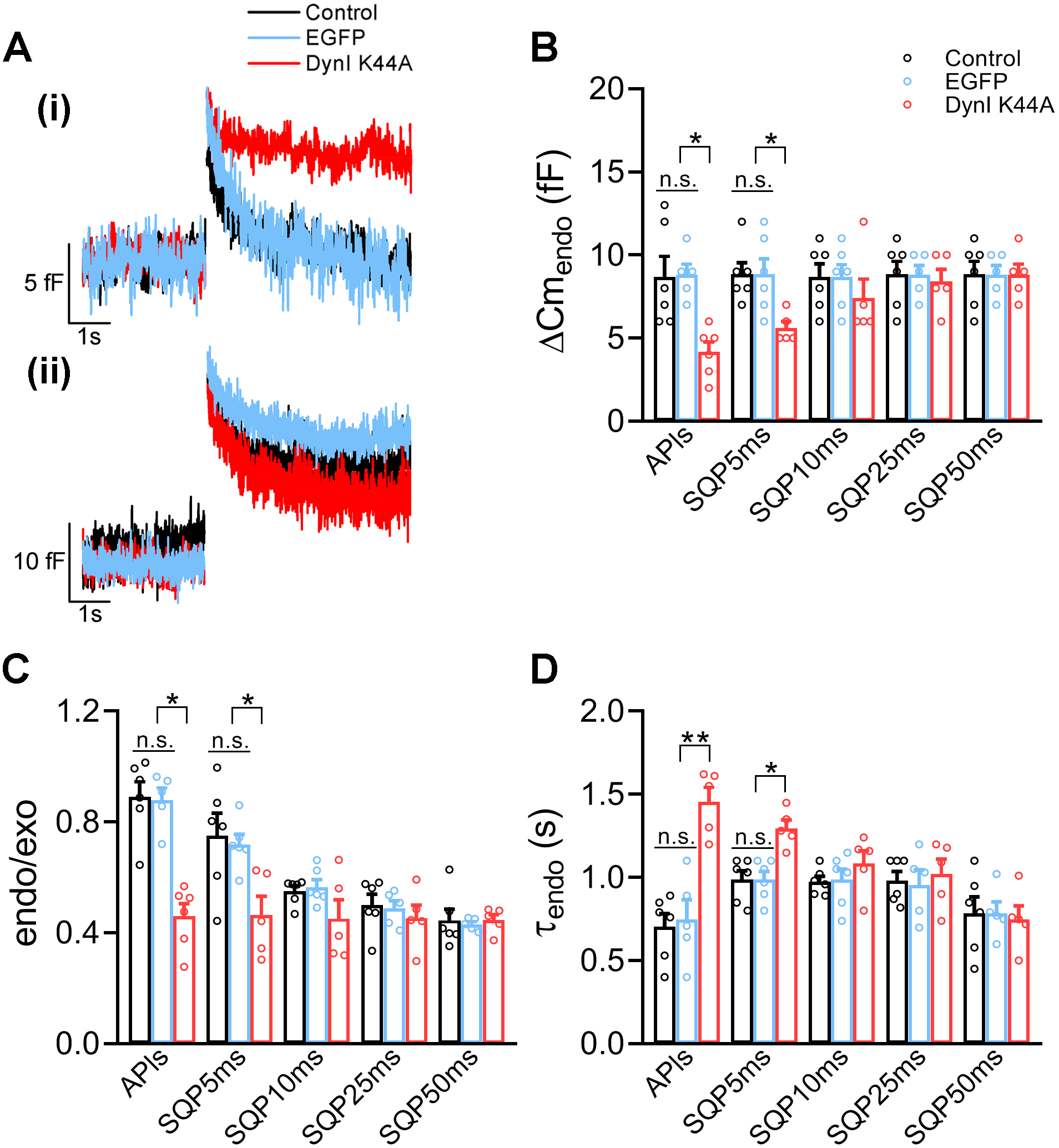
Expression of the dominant-negative mutant Dynamin-I K44A (DynI K44A) inhibits the endocytosis induced by APls and SQP5ms but not by longer stimuli: (A), Representative examples of Cm recordings obtained in response to the application of APls (i) and SQP50ms (ii) in a mock non-transfected cell (Control, black), in a cell expressing DynI K44A (red) and a cell expressing EGFP (blue). (B), (C) and (D), The plots represent average values, standard errors and individual measurements (one measurement/cell) of ΔCm_endo_, endo/exo ratio and τ_endo_ obtained by application of APls, SQP5ms, SQP10ms, SQP25ms and SQP50ms in Control (black), DynI K44A (red) and EGFP (blue). The data were analyzed by one-way ANOVA followed by Bonferroni comparisons. *p<0.05, **p<0.01. The sample sizes (number of individual cells) and the number of culture preparations from where these cells were obtained are summarized in Supplementary Table 1.

**Figure 7:**
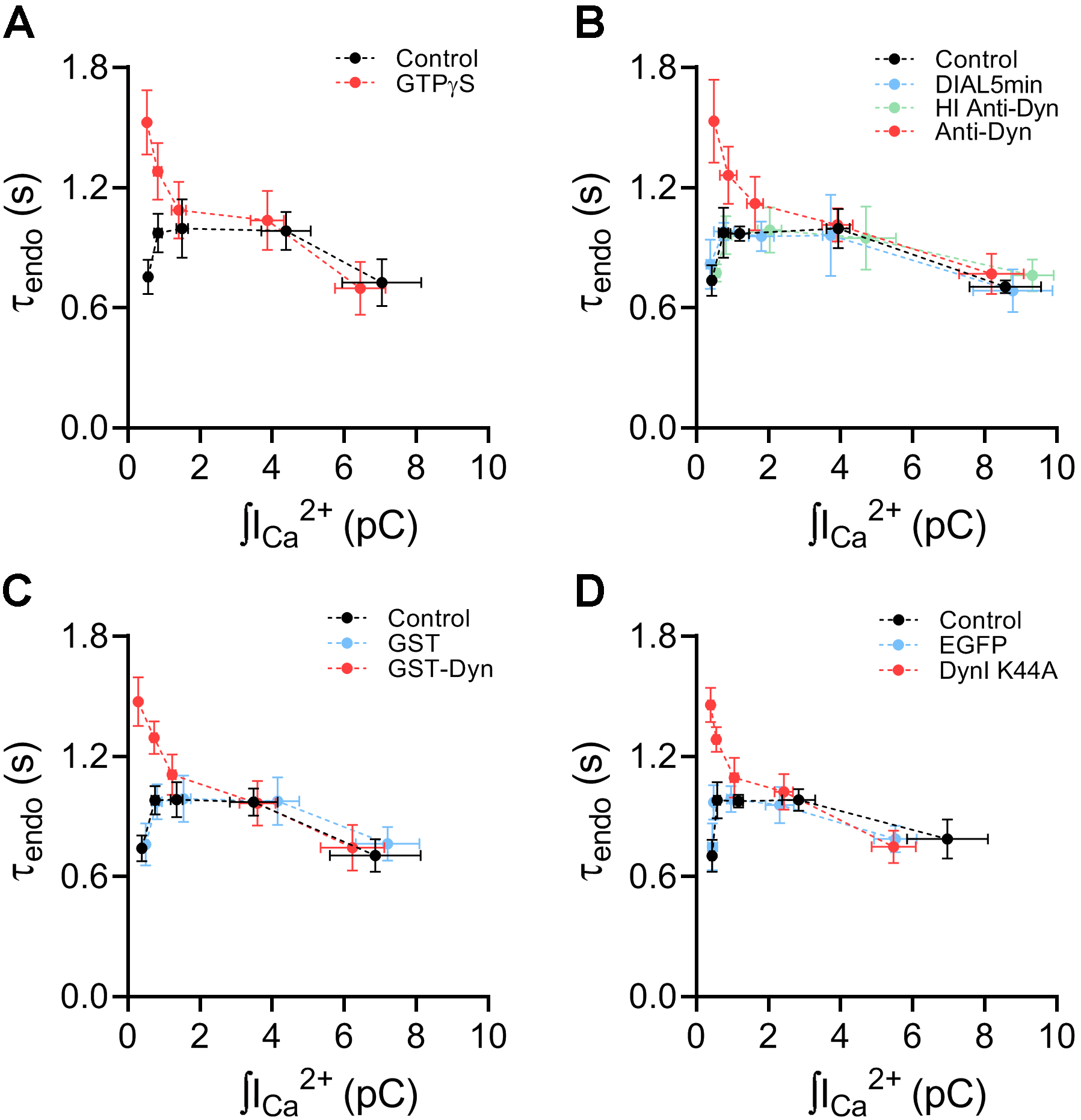
The endocytosis insensitive to dynamin inhibition is accelerated with the increase of Ca^2+^ entry. (A), (B), (C) and (D), The plots represent average values and standard errors of τ_endo_, in y-axis, versus ∫I_Ca2+_, in x-axis, obtained by application of APls, SQP5ms, SQP10ms, SQP25ms and SQP50ms in the different experimental conditions represented in Figures 3-6.

### The endocytosis activated after the complete IRP release is regulated by protein kinase C

Since the previous results of this work indicate that the dynamin-independent endocytosis activated after the total IRP exocytosis is Ca^2+^ dependent, we decided to evaluate the dependency of this process with protein kinase C (PKC). This decision was made because of the well-known Ca^2+^ sensitivity of this protein, as well as its well-documented participation in the regulation of endocytosis (Alvi *et al*. 2007; Jin *et al*. 2019; Chan & Smith 2003). We repeated the previously described stimuli protocol (i.e., APls, SQP5ms, SQP10ms, SQP25ms and SQP50ms) in presence of different experimental treatments that affect the activity of PKC (see Figures 8, 9, and 10, and Supplementary Figures 10, 11, and 13). First, we applied staurosporine (100 nM) in the external solution, which is very effective to block PKC (Tamaoki *et al*. 1986). Since staurosporine is a broad-spectrum kinase inhibitor, affecting other protein kinases (Ruegg & Burgess 1989; Wu-Zhang & Newton 2013), we performed similar experiments with bisindolylmaleimide XI (BIS XI, 3 μM), an inhibitor which is very specific for Ca^2+^-dependent PKC isoforms (Wilkinson *et al*. 1993; Wu-Zhang & Newton 2013). In both treatments, the drugs were added 5 min before starting with the measurements and maintained throughout the experiment. As a third approach to inhibit PKC, we applied phorbol-12-myristate-13-acetate (PMA, 4 μM) for 150 min at 37° C on cells kept in culture media with regulated CO_2_. It was previously described that high concentrations of PMA during long periods of time promote the migration of PKC to plasma membrane, its posterior ubiquitination and degradation (Krug *et al*. 1987; Lu *et al*. 1998). Finally, to increase PKC activity we applied PMA (100 nM) to external solution at room temperature, 7 min before and throughout the measurements. It is known that this PMA concentration stimulates PKC activity (Vitale *et al*. 1992; Wu-Zhang & Newton 2013). For all these treatments, we performed simultaneous control experiments with vehicle in absence of the drugs (for staurosporine and BIS XI we added the same amounts of DMSO (1/1000), and for PMA we added the same amounts of ethanol (EtOH, 1/1000), as were applied with the drugs). We did not obtain statistical differences in any of the measured parameters between both control conditions with ethanol (7 min at room temperature vs. 150 min at 37° C, see Supplementary Figure 12). In consequence, we grouped the control data for statistical comparisons against PMA data. In addition, none of the pharmacological treatments mentioned above affected I_Ca2+_ or ∫I_Ca2+_, and only 4 μM PMA modified (increased) ΔCm_exo_ particularly for SQP50ms (see examples of ΔCm recordings in Figures 8A, 9A, and 10A, and the summary of the data in the Supplementary Figures 10, 11, and 13). The effect of 4 μM PMA on exocytosis might be due to the previously reported action of this drug on vesicle mobilization (Vitale *et al*. 1995). Consistently with our hypothesis, staurosporine, BIS XI and 4 μM PMA significantly reduced the amplitudes of endocytosis (ΔCm_endo_) and endo/exo ratio for SQP25ms and SQP50ms, without affecting the same parameters for shorter stimuli (with the exception of 4 μM PMA, which also affected SQP10ms) (Figures 8B-C, 9B-C, and 10B-C). Additionally, all these treatments increased τ_endo_ (reduced the rate of endocytosis) for SQP25ms and SQP50ms, but they did not affect this parameter for the rest of stimuli (Figures 8D, 9D, and 10D). These results consistently indicate that the dynamin-independent endocytosis induced after SQP25ms and SQP50ms is PKC sensitive. It is surprising anyway that for these depolarizations, where most of IRP is released, membrane retrieval in our control conditions cannot compensate more than 50% of previous exocytosis (see endo/exo ratios along the work). However, the results obtained with 100 nM PMA (Figure 10) show that this mechanism has the potential capacity to retrieve a bigger amount of membrane. In fact, the application of this treatment induced a significant increase in the amplitude of endocytosis (ΔCm_endo_) and in the endo/exo ratio for SQP10ms, SQP25ms and SQP50ms (see Figure 10A(ii), B and C). In conclusion, the results of this and previous sections together indicate that when IRP is mostly released by SQP25ms or SQP50ms, membrane retrieval is mainly achieved by a dynamin-independent and Ca^2+^- and PKC-dependent endocytosis.

**Figure 8:**
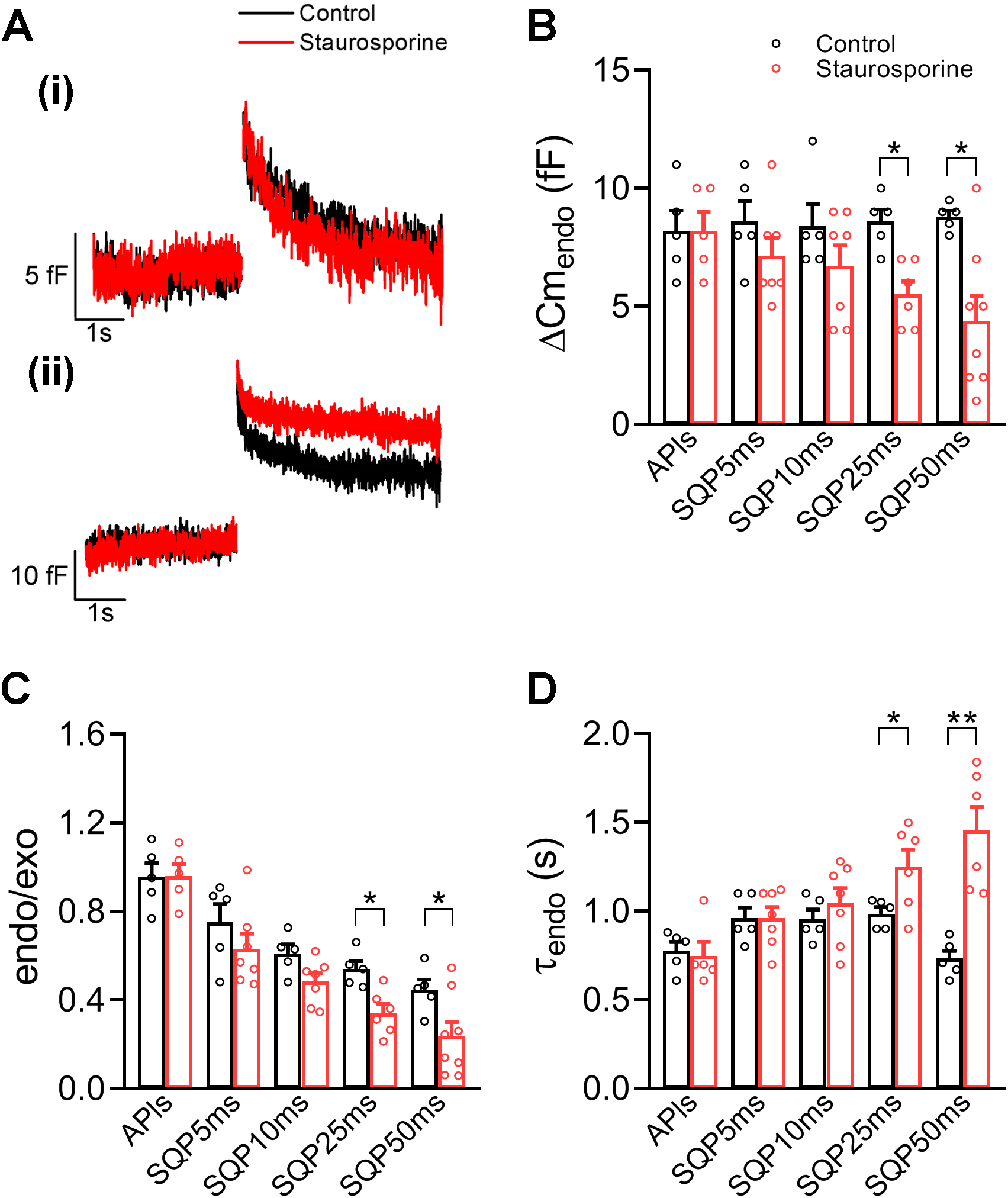
Application of protein kinase inhibitor staurosporine only inhibits the endocytosis induced by SQP25ms and SQP50ms: (A), Representative examples of Cm recordings obtained in response to the application of APls (i) and SQP50ms (ii) in two independent cells bathed in standard external solution (with addition of DMSO) (Control, black) or in presence of 100 nM staurosporine (red). (B), (C) and (D), The plots represent average values, standard errors and individual measurements (one measurement/cell) of ΔCm_endo_, endo/exo ratio and τ_endo_ obtained by application of APls, SQP5ms, SQP10ms, SQP25ms and SQP50ms in Control (black) and staurosporine (red). The data were analyzed by Student’s ‘t’ test. *p<0.05, **p<0.01. The sample sizes (number of individual cells) and the number of culture preparations from where these cells were obtained are summarized in Supplementary Table 1.

**Figure 9:**
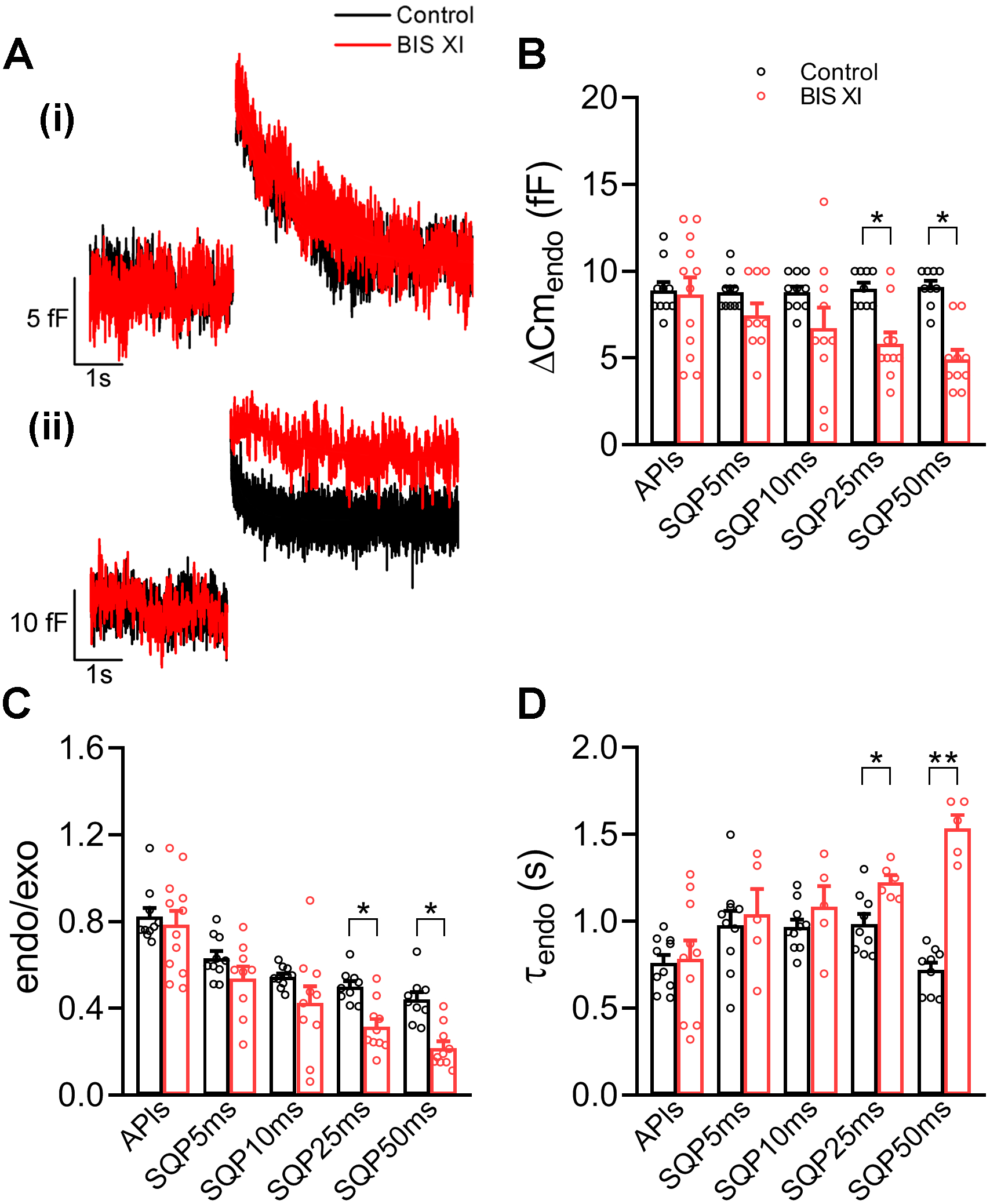
Application of protein kinase C inhibitor bisindolylmaleimide XI (BIS XI) only inhibits the endocytosis induced by SQP25ms and SQP50ms: (A), Representative examples of Cm recordings obtained in response to the application of APls (i) and SQP50ms (ii) in two independent cells bathed in standard external solution (with addition of DMSO) (Control, black) or in presence of 3 μM BIS XI (red). (B), (C) and (D), The plots represent average values, standard errors and individual measurements (one measurement/cell) of ΔCm_endo_, endo/exo ratio and τ_endo_ obtained by application of APls, SQP5ms, SQP10ms, SQP25ms and SQP50ms in Control (black, n≥9) and BIS XI (red, n≥9). The data were analyzed by Student’s ‘t’ test. *p<0.05, **p<0.01. The sample sizes (number of individual cells) and the number of culture preparations from where these cells were obtained are summarized in Supplementary Table 1.

**Figure 10:**
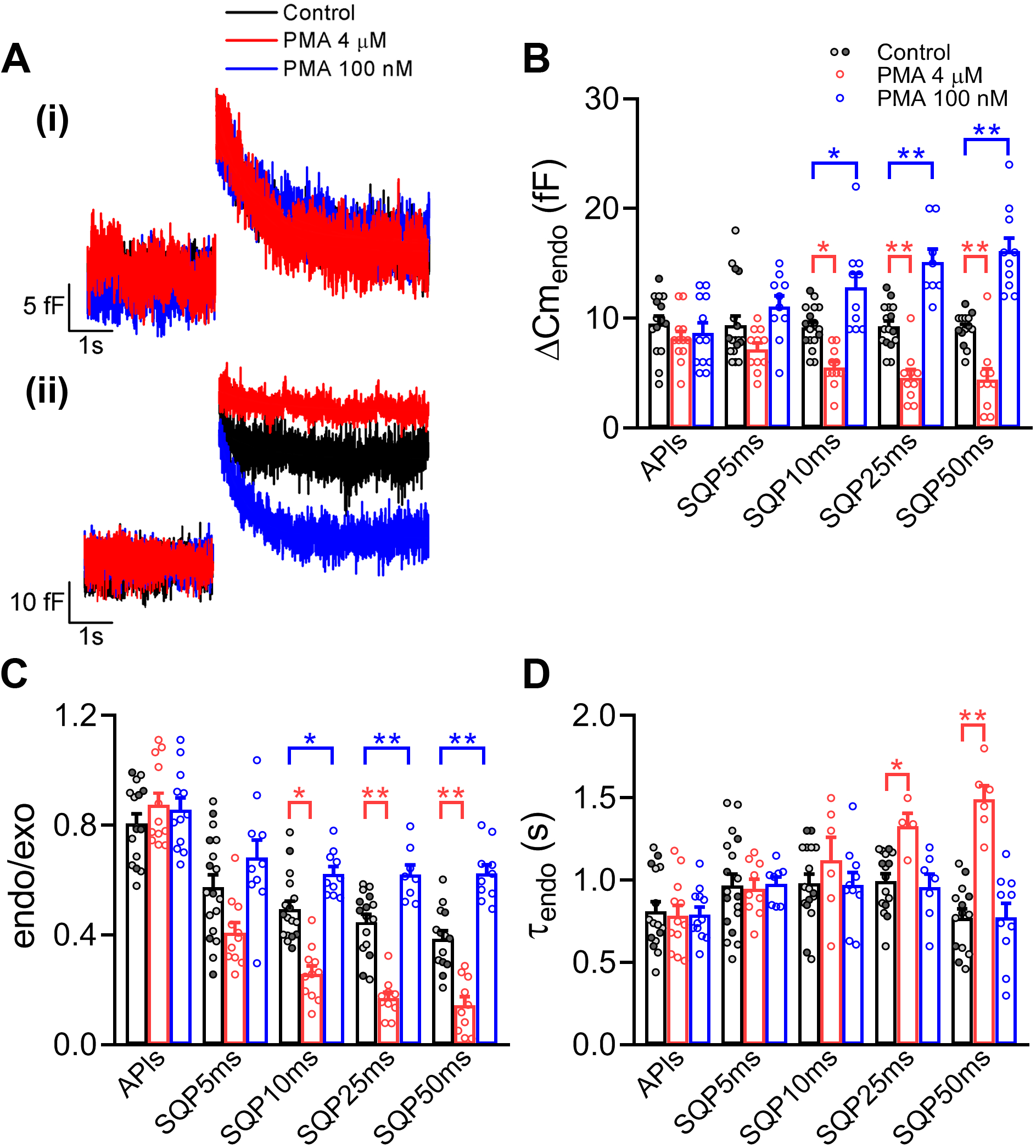
Application of phorbol esters modulates the endocytosis induced by SQP25ms and SQP50ms: (A), Representative examples of Cm recordings obtained in response to the application of APls (i) and SQP50ms (ii) in three independent cells bathed in standard external solution (with addition of ethanol) (Control, black), pretreated with 4 μM PMA during 150 min at 37° C (PMA 4 μM, red), or pretreated with 100 nM PMA during 7 min at room temperature (PMA 100 nM, blue). The Control Cm recording was obtained in a cell pretreated with ethanol during 7 min at room temperature. Control experiments with ethanol at both experimental conditions (7 min at room temperature and 150 min at 37° C) were not statistically different, in any measured parameter (Supplementary Figure 12). (B), (C) and (D), The plots represent average values, standard errors and individual measurements (one measurement/cell) of ΔCm_endo_, endo/exo ratio and τ_endo_ obtained by application of APls, SQP5ms, SQP10ms, SQP25ms and SQP50ms in Control (dark and light gray), 4 μM PMA (red), and 100 nM PMA (blue). Dark and light gray points represent individual Control measurements obtained with ethanol applied during 7 min at room temperature and during 150 min at 37° C, respectively. Control column bars and SE were obtained from the grouped Control data including both conditions, which was used for statistical comparisons against both PMA treatments. The data were analyzed by one-way ANOVA followed by Bonferroni comparisons. *p<0.05, **p<0.01. The sample sizes (number of individual cells) and the number of culture preparations from where these cells were obtained are summarized in Supplementary Table 1.

**Figure 11:**
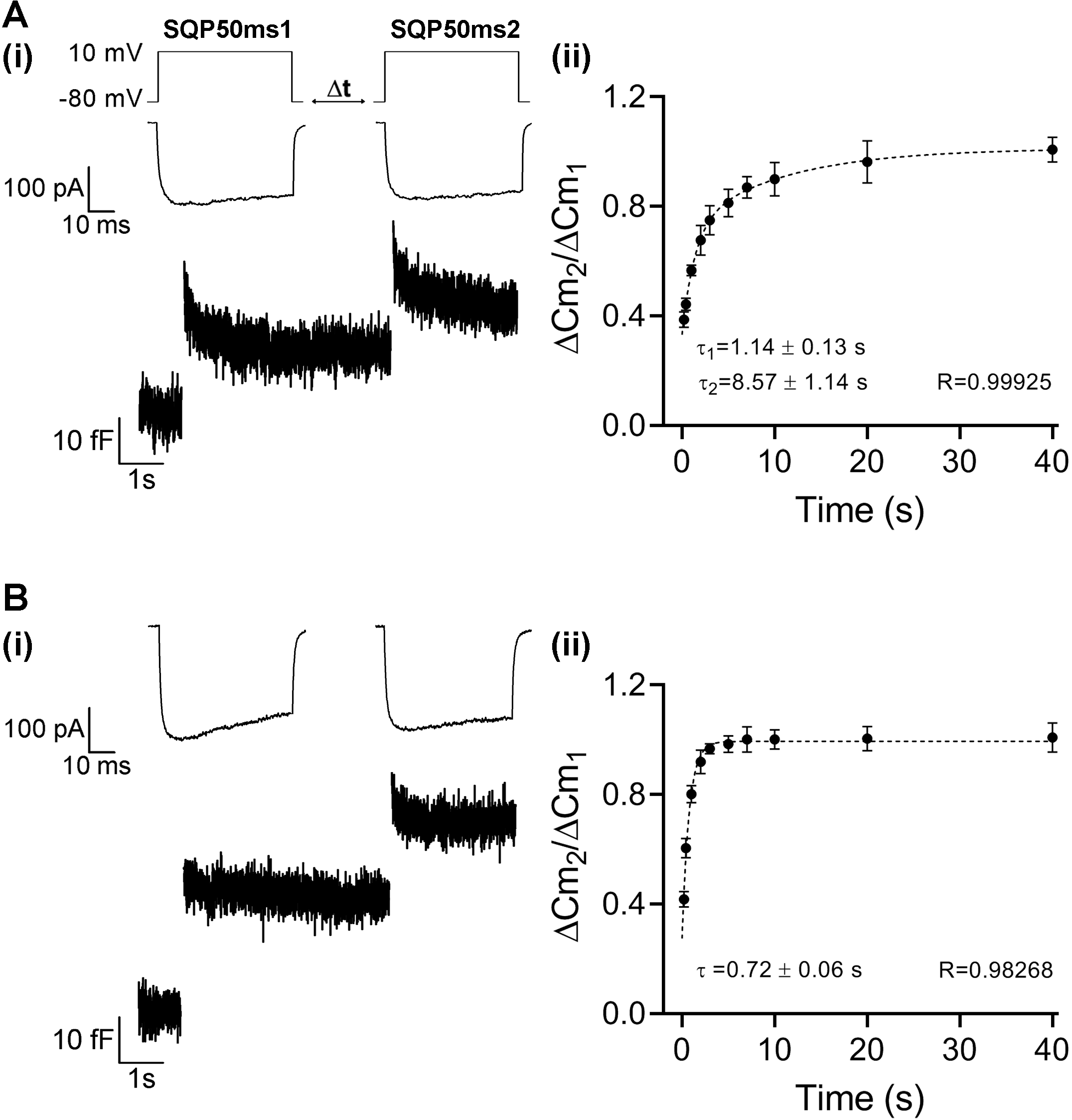
Protein kinase inhibitor staurosporine abolishes the slow component of IRP replenishment. In order to determine the kinetics of IRP replenishment, a pair of 50 ms depolarizations (SQP50ms_1_ and SQP50ms_2_, from −80 to +10 mV), with a variable time interval (Δt) between them, was applied in control conditions (A) and in presence of 100 nM staurosporine (B). The scheme of the stimulation protocol is represented at the top left of panel A. (i), I_Ca2+_ (above) and Cm (below) recordings obtained during a typical experiment when Δt = 5 s was applied. (ii), Relative replenishment of IRP (expressed as averages of ΔCm_2_/ΔCm_1_ for every Δt applied) was plotted against the Δt between paired SQP50ms pulses and fitted to a biexponential growing function of the form 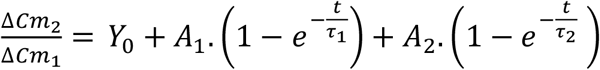 (A) or 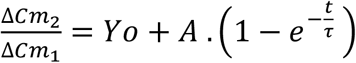 (B). The fitting parameters for the control condition (A) were Y_0_=0.32±0.01, A_1_=0.35±0.03, τ_1_=1.14±0.13 s, A_2_=0.33±0.03, and τ_2_=8.57±1.14 s, R^2^>0.9993; and for staurosporine (B) were Y_0_=0.25±0.03, A=0.73±0.03, and τ=0.72±0.06 s, R^2^>0.9827. The points represented in the plots are averages of all the measurements obtained in individual cells (one measurement per cell). The number of individual cells measured (between parentheses) were: for (A), 0.2 s (7), 0.4 s (7), 1 s (6), 2 s (6), 3 s (6), 5 s (7), 7 s (8), 10 s (7), 20 s (7) and 40 s (7); and for (B), 0.2 s (11), 0.4 s (11), 1 s (12), 2 s (12), 3 s (11), 5 s (11), 7 s (12), 10 s (10), 20 s (12) and 40 s (9), obtained in 9 independent cell culture preparations. The fittings were performed on average values.

### The inhibition of PKC abolishes the slow component of IRP replenishment

Previous results from our laboratory showed that after its complete exocytosis induced by SQP50ms, IRP recovers with two kinetically different components. While the fast component is dynamin- and F-actin dependent, and is negatively regulated by Ca^2+^, there is evidence that the slow component is positively regulated by cytoplasmic [Ca^2+^] (Montenegro *et al*. 2021). Therefore, we decided to analyze the dependence of IRP replenishment with PKC, a Ca^2+^-dependent protein kinase. To study the replenishment of IRP we applied a protocol composed of pairs of SQP50ms (Figure 11A(i), and Supplementary Figure 14), where the pulses of each pair are separated by time intervals (Δt) of 0.2, 0.4, 1, 2, 3, 5, 7, 10, 20 and 40 s (Montenegro *et al*. 2021). We expressed the degree of replenishment as the ratio between exocytosis evoked by the second stimulus (ΔCm_2_) and exocytosis evoked by the first stimulus (ΔCm_1_). Figure 11A(i) and B(i) show two typical examples of Ca^2+^ current (above) and Cm recordings (below) during the application of this protocol with a time interval of 5 s between stimuli (Δt= 5 s), in control conditions and in presence of staurosporine, respectively. A summary of I_Ca2+_, ∫I_Ca2+_, ΔCm_exo_, ΔCm_endo_, endo/exo and τ_endo_ of these experiments is represented in Supplementary Figure 15. The IRP replenishment (i.e., the averaged ΔCm_2_/ΔCm_1_ plotted versus the time interval between the first and the second stimulus) for these two conditions is represented in Figure 11A (ii) and B (ii), respectively. The averaged ΔCm_2_/ΔCm_1_ control data in Figure 11A (ii) were fitted to a double exponential growing function, obtaining a rapid time constant (τ_1_) of 1.14±0.13 s and a slow time constant (τ_2_) of 8.57±1.14 s (R^2^= 0.9993, reduced Chi-square=0.02621). The quality of this fitting was significantly better (F=100.2; p<0.05) than the one obtained by the application of a single exponential (R^2^=0.9779, reduced Chi-square=0.76878), as it was demonstrated by the application of a Fisher Test and in agreement with a previous publication from our group (Montenegro *et al*. 2021). On the other hand, the application of a double exponential to the averaged ΔCm_2_/ΔCm_1_ staurosporine data represented in Figure 11B (ii) (R^2^=0.9899, reduced Chi-square=0.26157) did not improve the quality of the fitting obtained with a single exponential growing function (R^2^=0.9827, reduced Chi-square=0.32184, F=38.1, p>0.2). It is important to mention that this last fitting provided a fast time constant τ of 0.72±0.06 s, indicating that staurosporine specifically inhibited the replenishment process associated with the slow component observed in control conditions.

As it was mentioned before, staurosporine is very efficient to inhibit PKC, but it is not specific for this protein kinase. Therefore, we also evaluated the IRP replenishment after the application of bisindolylmaleimide XI (BIS XI), with the same experimental protocol described before for staurosporine. Figure 12 represents examples of Ca^2+^ current (above) and Cm recordings (below) (i) obtained by the application of the paired pulse protocol with a time interval of 5 s between stimuli and IRP replenishment averaged plots (ii), for control conditions (A) and in presence of BIS XI (B). A summary of I_Ca2+_, ∫I_Ca2+_, ΔCm_exo_, ΔCm_endo_, endo/exo and τ_endo_ for these experiments is represented in Supplementary Figure 16. The replenishment results were very similar to the ones obtained with staurosporine: while the control averaged data points were better fitted to a double exponential growing function with a rapid and a slow component (for one exponential fitting: R^2^>0.9805 and reduced Chi-squared=0.38281; for the double exponential fitting: R^2^>0.9987 and reduced Chi-squared=0.02635; F=48.328; p<0.05), the averaged data points obtained in presence of BIS XI fitted very well to a single exponential growing function (R^2^>0.9861 and reduced Chi Squared=0.38714) and a double exponential function could not improve the quality of the fitting (R^2^> 0.9908 and reduced Chi-squared=0.14368; F=6.93564; p>0.3). This last single exponential fitting provided a fast time constant τ of 0.96±0.07 s, indicating that, as observed with staurosporine, also BIS XI specifically inhibited the replenishment process associated with the slow component of control conditions. The results obtained with staurosporine and BIS XI indicate that a PKC dependent step is specifically related to the regulation of the slow replenishment process observed in control conditions. Therefore, the results of this work indicate that PKC regulates the endocytosis and the replenishment produced after complete release of IRP by SQP50ms.

**Figure 12:**
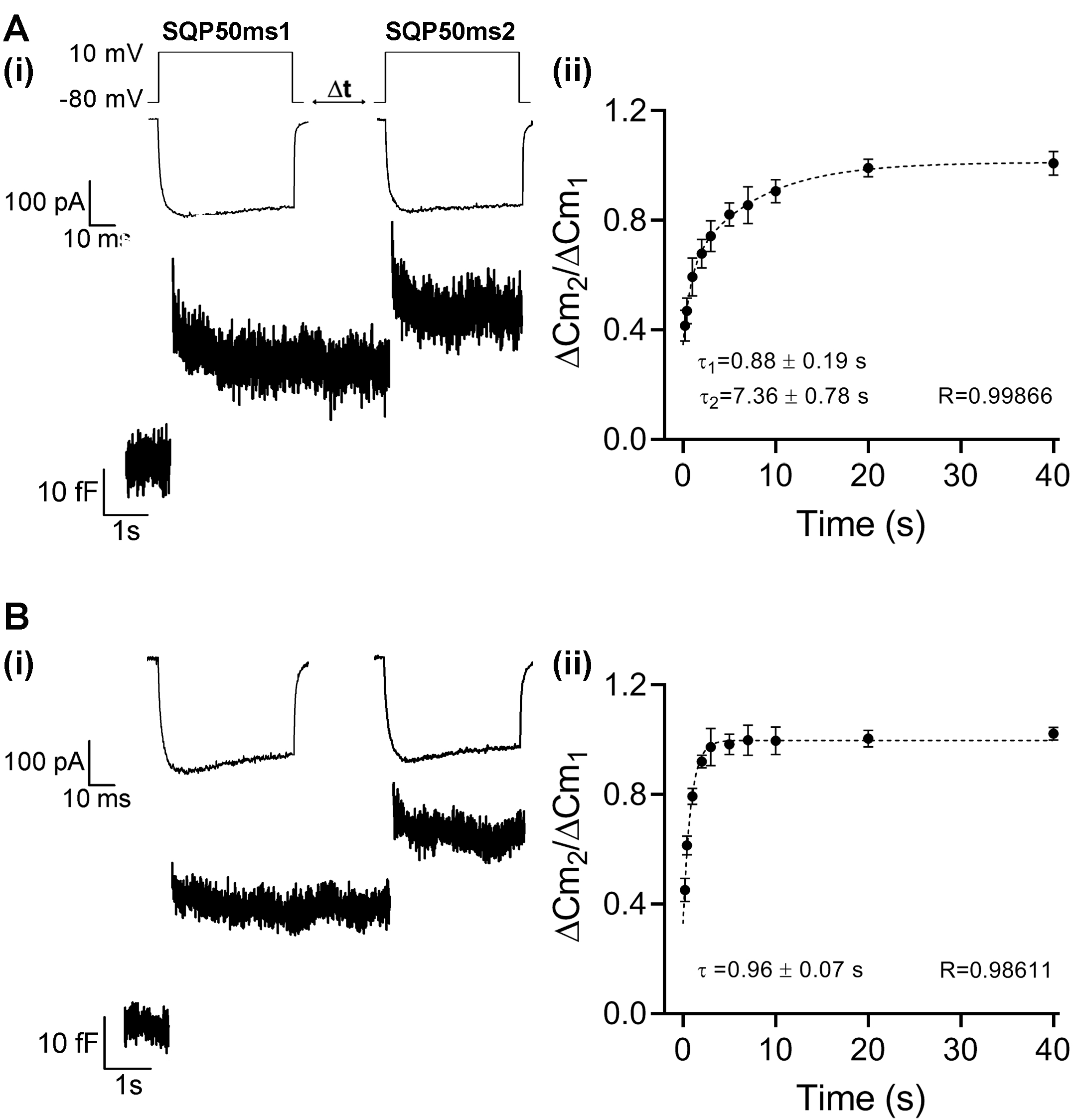
Protein kinase C inhibitor bisindolylmaleimide XI (BIS XI) abolishes the slow component of IRP replenishment. In order to determine the effect of PKC inhibition on the kinetics of IRP replenishment, a pair of 50 ms depolarizations (SQP50ms_1_ and SQP50ms_2_, from −80 to +10 mV) with variable time intervals (Δt) between them was applied in control conditions (A) and in presence of 3 μM BIS XI (B). The scheme of the stimulation protocol is represented at the top left of panel A. (i), I_Ca2+_ (above) and Cm (below) recordings obtained during a typical experiment when a Δt = 5 s was applied. (ii) Relative replenishment of IRP (expressed as averages of ΔCm_2_/ΔCm_1_ for every Δt applied) was plotted against the Δt between paired SQP50ms pulses and fitted to a biexponential growing function of the form 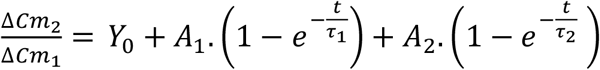 (A) or 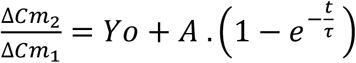 (B). The fitting parameters for the control condition were Y_0_=0.35±0.02, A_1_=0.39±0.03, τ_1_=0.88±0.19 s, A_2_=0.27±0.03, and τ_2_=7.36±0.78 s, R^2^>0.9987, and for BIS XI were Y_0_=0.37±0.03, A=0.63±0.03, and τ=0.96±0.07 s, R^2^>0.9861. The points represented in the plots are averages of all the measurements obtained in individual cells (one measurement per cell). The number of individual cells measured (between parentheses) were: for (A), 0.2 s (10), 0.4 s (11), 1 s (11), 2 s (10), 3 s (10), 5 s (11), 7 s (9), 10 s (9), 20 s (8) and 40 s (10); and for (B), 0.2 s (8), 0.4 s (9), 1 s (8), 2 s (8), 3 s (8), 5 s (9), 7 s (10), 10 s (9), 20 s (8) and 40 s (7), obtained in 9 independent cell culture preparations. The fittings were performed on average values.

## Discussion

The present study demonstrates that two rapid endocytic mechanisms contribute to the membrane retrieval produced after the exocytosis of the immediately releasable pool in mouse chromaffin cells. Each mechanism predominates in a different range of stimulus strength, or in other words, for a different fraction of released IRP. For brief stimuli, that release less than 50% of IRP, the dominant mechanism is a fast dynamin-dependent endocytosis. On the other hand, for more prolonged stimuli releasing bigger fractions of IRP, the prevalent mechanism is a fast dynamin-independent but Ca^2+^ entry- and PKC-dependent endocytosis.

In a previous work, we described a fast dynamin-dependent endocytosis that occurred after the application of APls (Moya-Diaz *et al*. 2016). In a more recent article, we demonstrated that this mechanism is also dependent on cortical F-actin (Montenegro *et al*. 2021). In the present work, we showed that this endocytic mechanism is severely attenuated after the application of depolarizations longer than APls and SQP5ms. We do not have a definitive answer for this attenuation. A similar phenomenon was observed after the application of increasing frequencies of action potential like stimuli in bovine chromaffin cells (Chan & Smith 2001). In neuron terminals, it was proposed that deceleration of endocytosis with the increase in stimulation might be due to saturation of the endocytic machinery (Sankaranarayanan & Ryan 2000), to accumulation of unretrieved vesicles at the plasma membrane (Sun *et al*. 2002), or to an inhibitory effect of cytosolic Ca^2+^ (von Gersdorff & Matthews 1994). Recently, we also showed in mouse chromaffin cells that an endocytic process, with similar characteristics than the one observed in the present work, is reduced and decelerated with the increased in basal Ca^2+^ concentration (Montenegro *et al*. 2021). We believe that the previously proposed mechanisms are not excluding, and therefore it is possible that more than one can contribute to the reduction of the fast dynamin-dependent endocytosis with the increase in pulse duration.

Different dynamin-independent endocytosis mechanisms were determined in non-neural cells, which are mostly associated with constitutive protein internalization, fluid phase endocytosis and regulation of membrane tension (Krag *et al*. 2010; Thottacherry *et al*. 2018; Bonazzi *et al*. 2005). Additionally, direct determinations of dynamin-independent endocytosis were obtained in cone photoreceptors of salamander retina (Van Hook & Thoreson 2012) and dorsal root ganglion neurons (Zhang *et al*. 2004), although in the latter the endocytosis was observed after a particular mechanism of Ca^2+^-independent exocytosis. To our knowledge, our work is the first direct demonstration of a dynamin-independent endocytosis activated after Ca^2+^-dependent exocytosis in neuroendocrine cells. Moreover, our results are consistent with previous indirect observations in chromaffin cells from other laboratories. For example, Chan et al described a calcineurin-independent endocytosis in bovine chromaffin cells stimulated with action potentials applied at low frequencies (Chan & Smith 2001). Even though calcineurin is a natural activator of dynamin, this phosphatase also modulates other proteins that may regulate endocytosis, such as L-type Ca^2+^ channels, amphiphysin, synaptojanin, etc (Cousin & Robinson 2001; Li *et al*. 2011). Additionally, Burgoyne group (Graham *et al*. 2002) also suggested the presence of a dynamin-independent membrane retrieval mechanism from amperometric results obtained in bovine chromaffin cells. Interestingly, they observed that PKC activation by PMA induced a reduction of amperometric spikes half-width, rise time and fall time in cells previously treated with dynamin inhibitors. Their interpretation was that PKC activation promotes a dynamin-independent fast kiss- and-run mode of fusion of secretory vesicles.

We demonstrated the presence of dynamin-independent endocytosis by application of four different experimental approaches that inhibit the action of dynamin, as was reported by previous studies. In different experimental series, we applied the GTP non-hydrolysable analogue GTPγS (Xu *et al*. 2008), the monoclonal antibody Anti-Dyn (Gonzalez-Jamett *et al*. 2010; Montenegro *et al*. 2021; Moya-Diaz *et al*. 2016; Moya-Diaz *et al*. 2020), the peptide GST-Dyn (Shupliakov *et al*. 1997; Fulop *et al*. 2008; Moya-Diaz *et al*. 2016), and the GTPase-defective dominant-negative mutant DynI-K44A (McMahon & Boucrot 2011). These four treatments revealed the presence of a dynamin-independent endocytosis that dominates membrane retrieval after the application of depolarizations longer than 10 ms, which induces the exocytosis of more than 50% of IRP.

A variety of results shown in this work indicate that the dynamin-independent endocytosis mechanism is accelerated by Ca^2+^ entry. First, the time constant τ_endo_ is significantly decreased when the duration of the stimulus is prolonged from SQP25ms to SQP50ms (Figure 1E). Second, in all the conditions where dynamin-dependent endocytosis is inhibited, τ_endo_ of the residual endocytosis process is consistently reduced with the size of ∫I_Ca2+_ (Figure 7). Third, τ_endo_ of the endocytosis activated after SQP50ms is significantly reduced with the increase in ∫I_Ca2+_ provoked by the augment of external [Ca^2+^] (Figure 2). Several authors reported a positive Ca^2+^ dependence for different types of endocytosis (Wu *et al*. 2009). The calcineurin-dependent compensatory endocytosis in bovine chromaffin cells is accelerated by the increase in Ca^2+^ entry (Engisch & Nowycky 1998; Smith & Neher 1997). Clathrin-dependent endocytosis is also positively modulated by Ca^2+^ in the calyx of Held and in mouse chromaffin cells (Hosoi *et al*. 2009; Yao *et al*. 2012). Moreover, Ca^2+^ influx initiates bulk endocytosis in the calyx of Held (Wu *et al*. 2009). Finally, a threshold level of Ca^2+^ entry has to be surpassed to trigger excess retrieval (Smith & Neher 1997; Perez Bay *et al*. 2012). Our results suggest that at least part of the positive Ca^2+^ dependence observed in the dynamin-independent endocytosis described in this work can be explained by a PKC-dependent regulation of this process. We demonstrated this PKC dependency by evaluating the effect of the inhibition of this enzyme with three independent experimental treatments, i.e., the non-specific kinase inhibitor staurosporine (Figure 8), the very specific PKC inhibitor bisindolylmaleimide XI (Figure 9), and a prolonged treatment with high concentration of PMA (Figure 10), which provokes the degradation of PKC (Krug *et al*. 1987). Additionally, the application of 100 nM PMA during a short period of time, which is a well-known stimulator of PKC (Vitale *et al*. 1992), promotes a significant increase in the dynamin-independent endocytosis. This last result also indicates that this fast dynamin-independent endocytosis can potentially be up-regulated, possibly overcoming the incomplete compensation of exocytosis observed in our control experiments.

Other investigators described that PKC is also involved in the regulation of vesicle mobilization to plasma membrane, and consequently in vesicle pools replenishment (Vitale *et al*. 1992; Trifaro *et al*. 2008). Our results show that PKC participates in the regulation of the slow component of IRP replenishment. Therefore, it seems that PKC may play a pivotal role in the recycling of secretory vesicles associated with IRP exocytosis, regulating dynamin-independent endocytosis and replenishment of this pool.

It is believed that IRP can be important in secretion during action potentials firing at low basal frequencies (Moya-Diaz *et al*. 2016; Oré & Artalejo 2005). Previous results from our laboratory show a central role of dynamin in the control of fast endocytosis and rapid replenishment after ETAP (exocytosis triggered by action potential) exocytosis, a fraction of IRP released by APls (Moya-Diaz *et al*. 2016). The present work adds information on other mechanisms of endocytosis and replenishment operating when bigger fractions of IRP are released. Dynamin-dependent and dynamin-independent mechanisms operating together may be essential for IRP sustainability during repetitive stimulation.

## Abbreviations

Anti-Dyn: monoclonal antibody against dynamin I & II
Anti-Dyn HI: Heat Inactivated Anti-Dyn
APls: action potential-like stimulus
BIS XI: bisindolylmaleimide XI
[Ca^2+^]_o_: Ca^2+^ concentration of the external solution
Cm: cell membrane capacitance
ΔCm_exo_: synchronous increase in Cm
ΔCm_endo_: decrease in Cm
DIAL5m: dialysis with standard internal solution for 5 min
DMSO: dimethylsulfoxide
Dynamin-I K44A: a dynamin-I GTPase-defective dominant-negative mutant
EGFP: Enhanced Green Fluorescent Protein
endo/exo: ratio between ΔCm_endo_ and ΔCm_exo_
ETAP: exocytosis triggered by action potential
EtOH: ethanol
F-actin: filamentous actin
GST: glutathione S-transferase
GST-Dyn: GST-Dyn829-842, peptide containing the recognition motif for SH3 domain-containing proteins of the dynamin-1 proline-rich domain attached to GST
GTPγS: non-hydrolyzable GTP analogue
I_Ca2+_: amplitude of Ca^2+^ currents
∫I_Ca2+_: integral of Ca^2+^ currents
L-type Ca^2+^: L-type calcium channels
IRP: immediately releasable pool
PKC: protein kinase C
PMA: phorbol-12-myristate-13-acetate
SQP: square depolarization pulse
τ_endo_: time constant of endocytosis
VDCC: voltage dependent Ca^2+^ channels.

## Acknowledgments

This work was supported by the grants PICT 0524-2014 and PICT 2764-2016 from the Agencia Nacional de Promoción Científica y Tecnológica, and UBACyT 2014-2017 from the Universidad de Buenos Aires.

