## Supplementary Material for "Membrane Retrieval after Immediately Releasable Pool (IRP) Exocytosis is produced by Dynamin-Dependent and Dynamin-Independent Mechanisms"

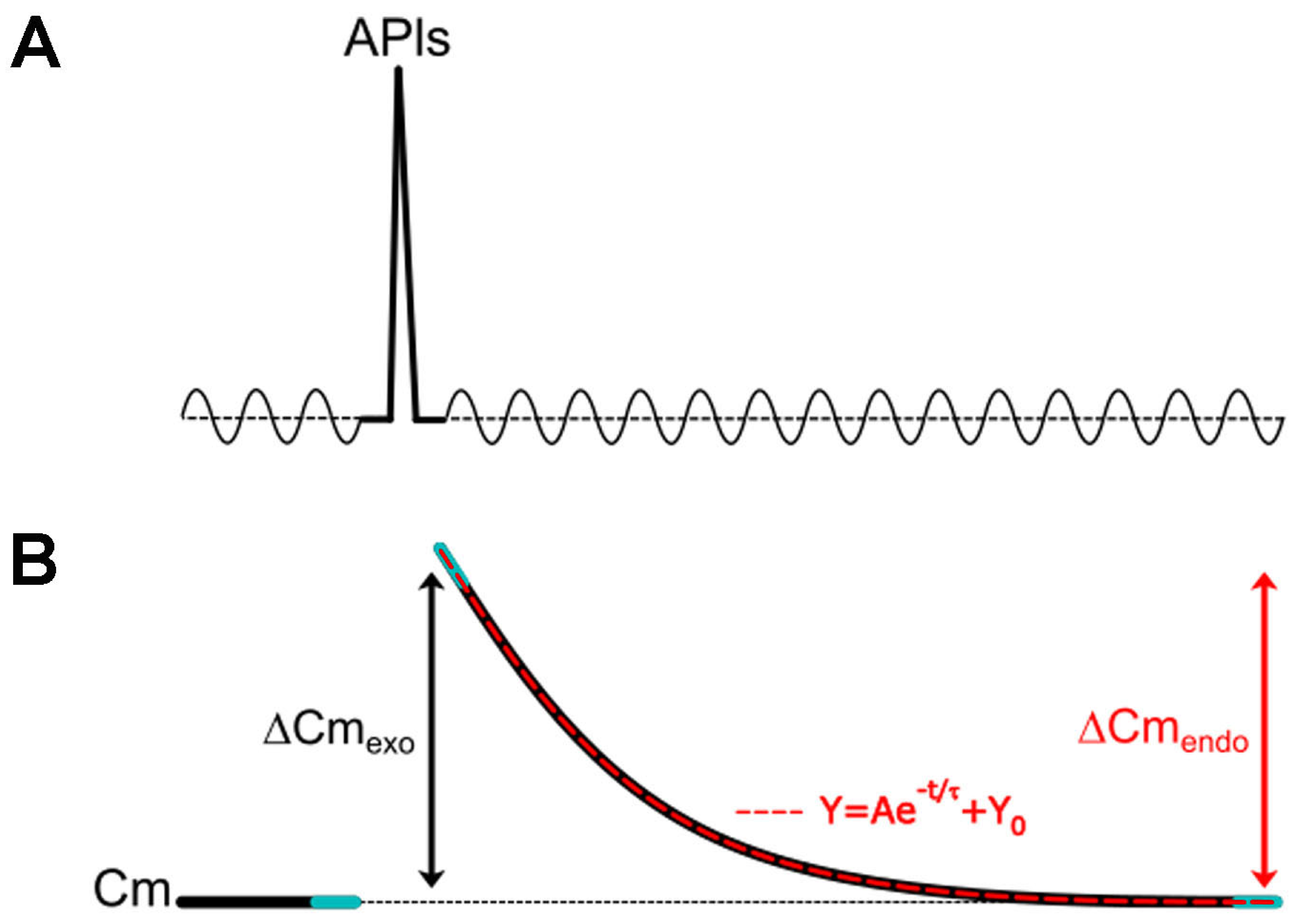


**Supplementary Figure 1:** Scheme representing the change in capacitance induced by a voltage pulse (in this case to APIs, but it can be applied to any type of pulse used in this work). The light blue lines represent the portions of the recording used to estimate the Cm values at the initial baseline, at the peak of Cm increase after the pulse and at the end of Cm decay, from where $\Delta Cm_{exo}$ and $\Delta Cm_{endo}$ were estimated. The dotted red line represents the portion of the recording used for the single exponential fitting of endocytosis.

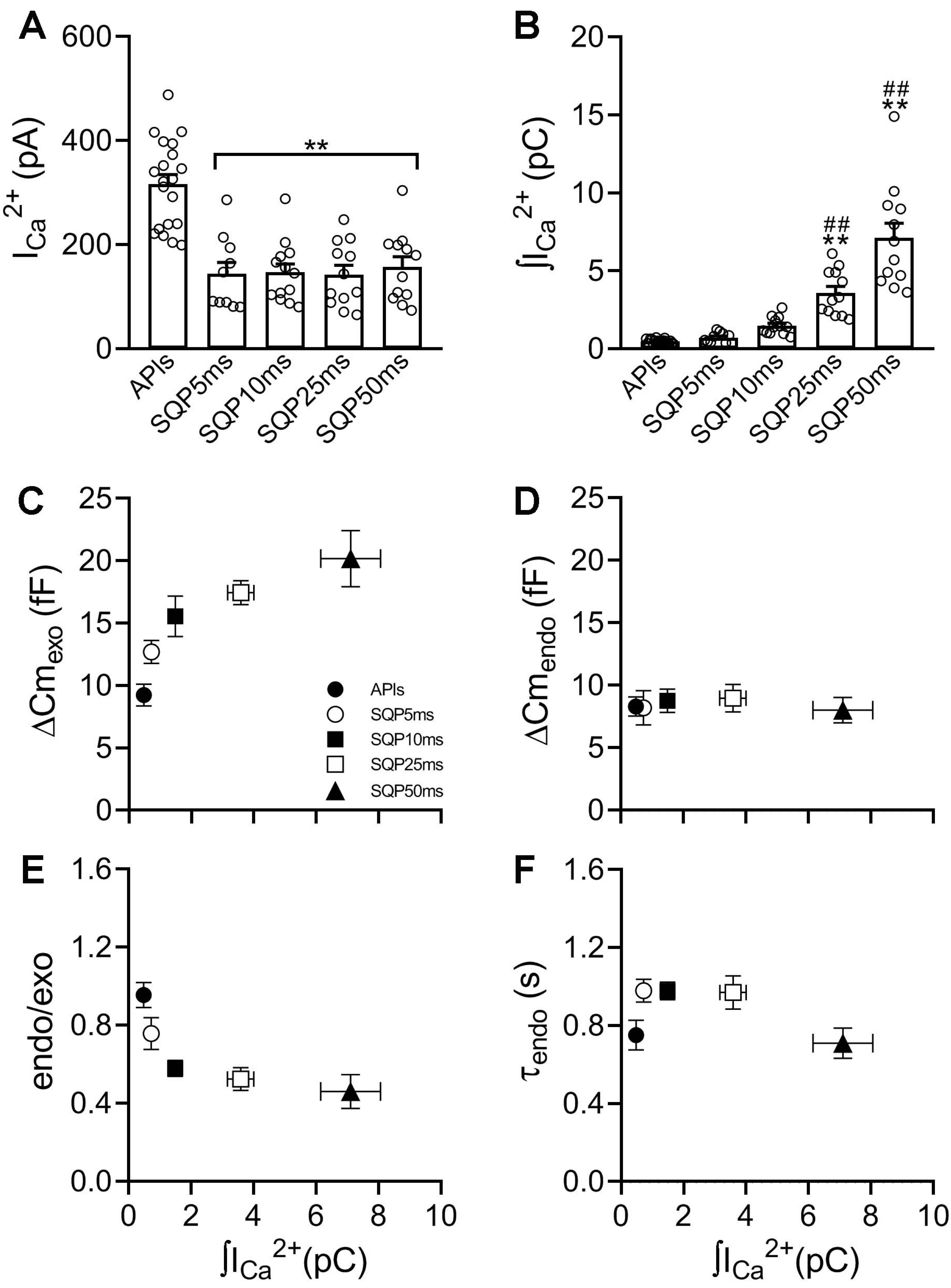


**Supplementary Figure 2:** (A) and (B), The plots represent average values, standard errors and individual measurements (one measurement/cell) of $I_{Ca}2+$ and $\int I_{Ca}2+$ obtained by application of APls, SQP5ms, SQP10ms, SQP25ms and SQP50ms. The data were analyzed by one-way ANOVA followed by Bonferroni comparisons. ** or ## represent $p<0.01$, where * symbols represent comparison of every condition versus APls, and # symbols represent comparison of every condition versus SQP5ms. (C), (D), (E) and (F), The plots represent average values and standard errors of $\Delta Cm_{exo}$, $\Delta Cm_{endo}$, endo/exo ratio and $\tau_{endo,}$ in y-axis, versus $\int I_{Ca}2+$, in x-axis, obtained by application of APls, SQP5ms, SQP10ms, SQP25ms and SQP50ms. The sample sizes (number of individual cells) and the number of culture preparations from where these cells were obtained are summarized in Supplementary Table 1. The sample sizes (number of individual cells) and the number of culture preparations from where these cells were obtained are summarized in the Supplementary Table 1 (first line, Control).

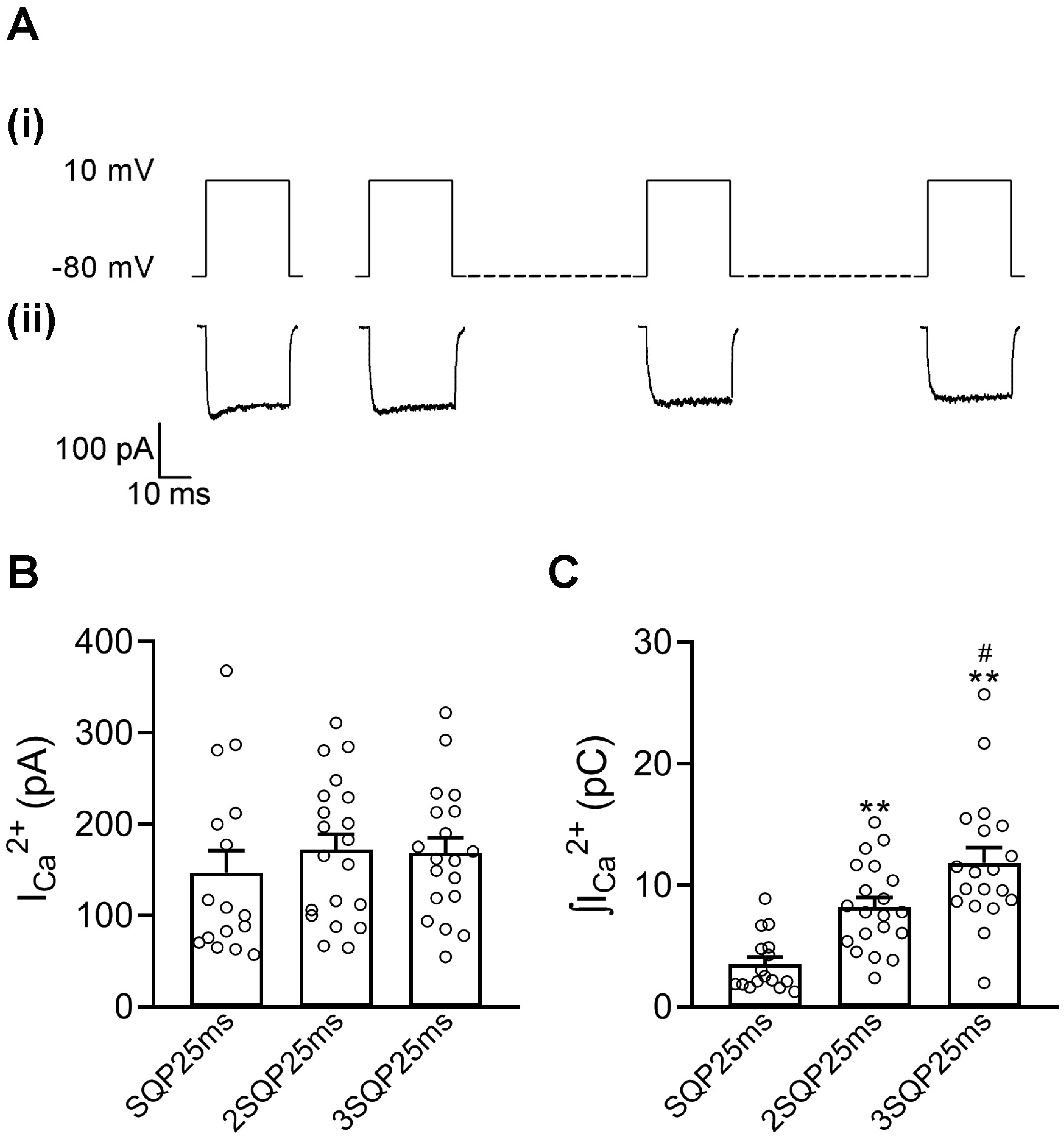


**Supplementary Figure 3:** (A) Representation of one (left) and three (right) consecutive SQP25ms at 13 Hz (i), and typical examples of $Ca^{2+}$ currents (ii) obtained in response to the application of these stimuli. (B) and (C), The plots represent average values, standard errors and individual measurements (one measurement/cell) of $I_{Ca}2+$ and $\int I_{Ca}2+$ obtained by application of one (n=16), two (n=20) or three (n=19) consecutive SQP25ms at 13 Hz, which were obtained in 16 independent cell culture preparations. The data were analyzed by one-way ANOVA followed by Bonferroni comparisons. # $p < 0.05$, ** $p < 0.01$. * symbols represent comparison of every condition versus SQP25ms, and # symbols represent comparison between 2SQP25ms and 3SQP25ms.

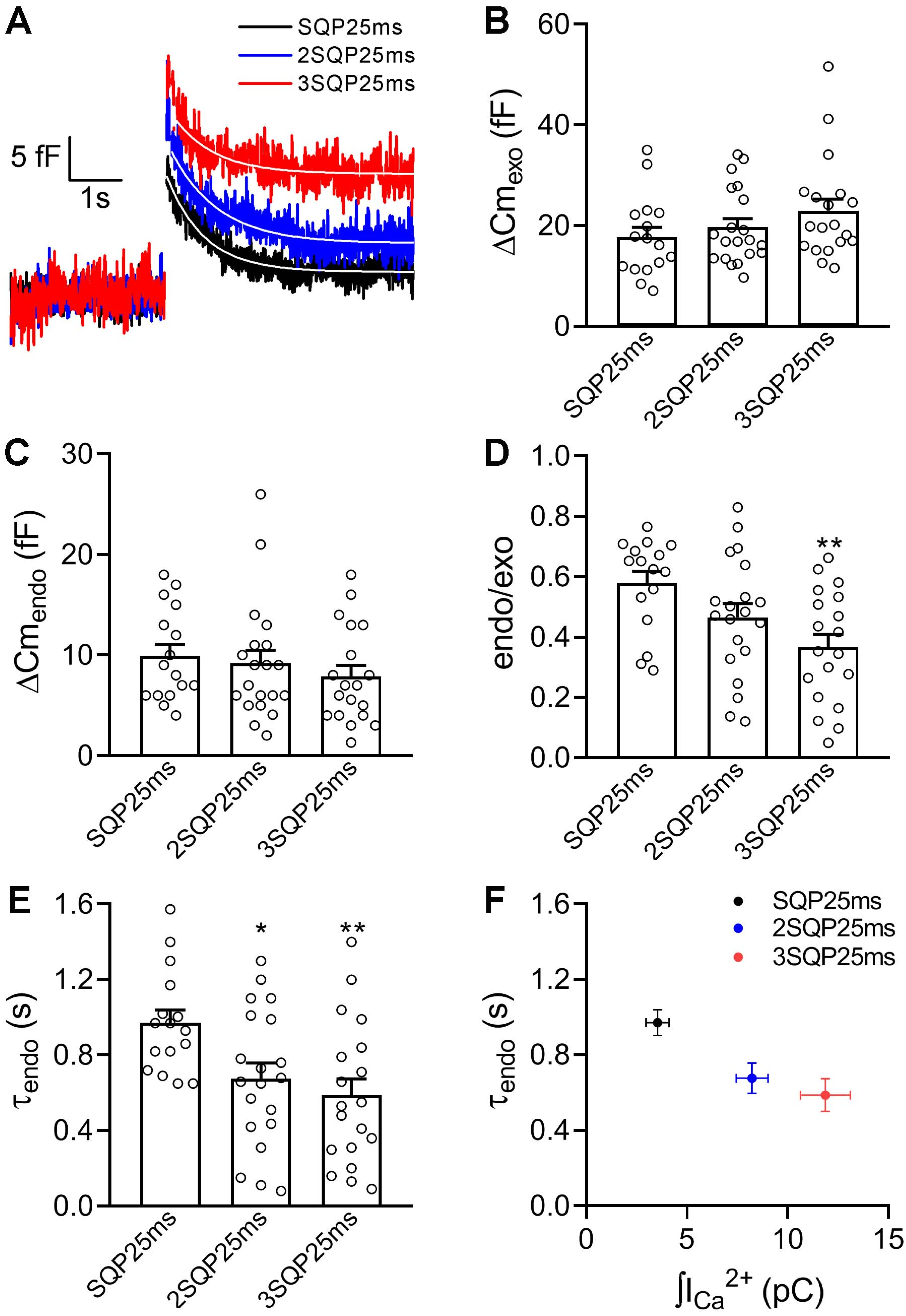


**Supplementary Figure 4:** (A) Typical examples of Cm recordings obtained in response to the application of one, two or three consecutive SQP25ms at 13 Hz. The white lines represent single exponential fittings ($\Delta Cm = Yo + A.e^{-\tau endo}$) applied to the decay in Cm, where $\tau_{endo}$ was 0.92 s ($R^2 > 0.8196$), 0.79 s ($R^2 > 0.8783$) and 0.59 s ($R^2 > 0.4979$) for SQP25ms, 2SQP25ms and 3SQP25ms, respectively. (B), (C), (D) and (E), The plots represent average values, standard errors and individual measurements (one measurement/cell) of $\Delta Cm_{exo}$, $\Delta Cm_{endo}$, endo/exo ratio and $\tau_{endo}$ obtained by application of the three types of stimuli described in (A) (n=16, 20 and 19, respectively, obtained in 16 independent cell culture preparations). The data were analyzed by one-way ANOVA followed by Bonferroni comparisons. * $p < 0.05$, ** $p < 0.01$, representing comparisons versus one SQP25ms. (F), Averaged values of $\tau_{endo}$ plotted against averaged $\int I_{Ca^{2+}}$.

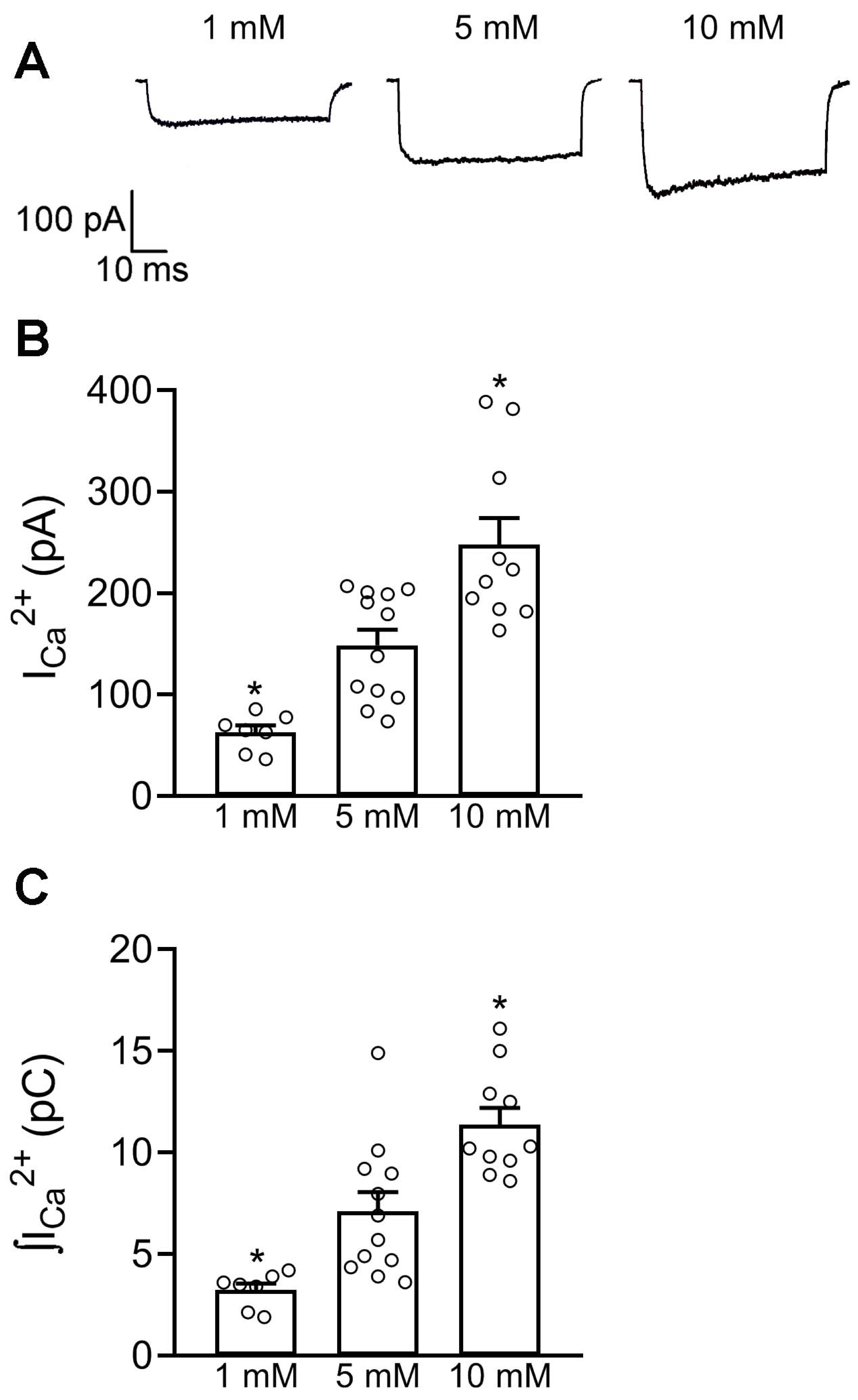


**Supplementary Figure 5:** (A), Typical examples of $Ca^{2+}$ currents obtained in response to the application of SQP50ms in presence of 1, 5 and 10 mM $[Ca^{2+}]_o$ in three independent cells. (B) and (C), The plots represent average values, standard errors and individual measurements (one measurement/cell) of $I_{Ca}2+$ and $\int I_{Ca}2+$ obtained in the three conditions described in (A). The data were analyzed by one-way ANOVA followed by Bonferroni comparisons. * $p < 0.05$. The comparisons were done versus 5 mM $[Ca^{2+}]_o$. The sample sizes (number of individual cells) and the number of cell culture preparations from where these cells were obtained are summarized in the Supplementary Table 1.

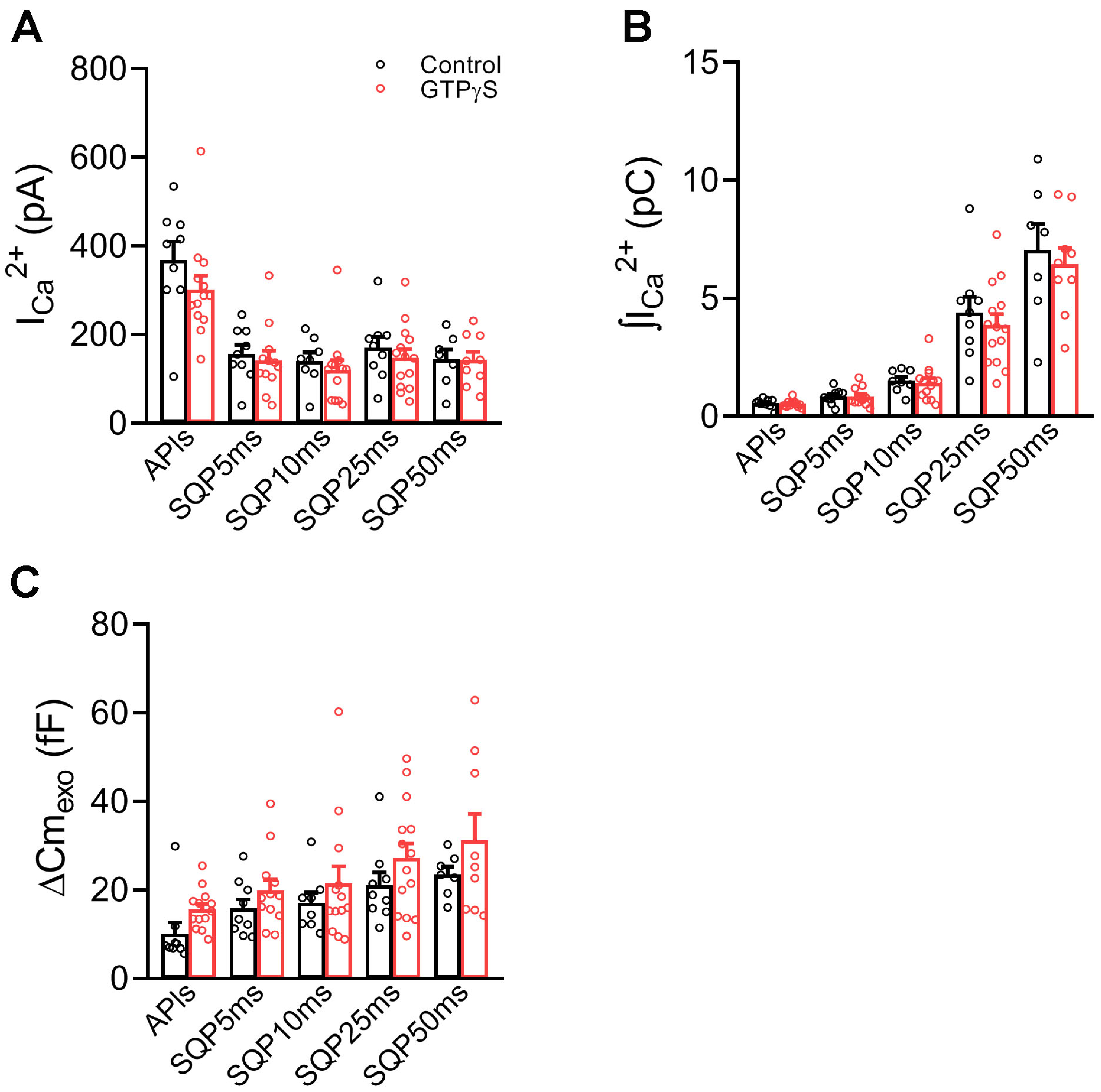


**Supplementary Figure 6:** (A), (B) and (C), The plots represent average values, standard errors and individual measurements (one measurement/cell) of $I_{Ca}2+$, $\int I_{Ca}2+$ and $\Delta Cm_{exo}$ obtained by application of APls, SQP5ms, SQP10ms, SQP25ms and SQP50ms in control conditions (standard internal solution, Control, black) or in presence of 0.3 mM GTPγS (red) in the internal solution, respectively. The data were analyzed by Student's 't' test. No difference was found between conditions. The sample sizes (number of individual cells) and the number of cell culture preparations from where these cells were obtained are summarized in the Supplementary Table 1.

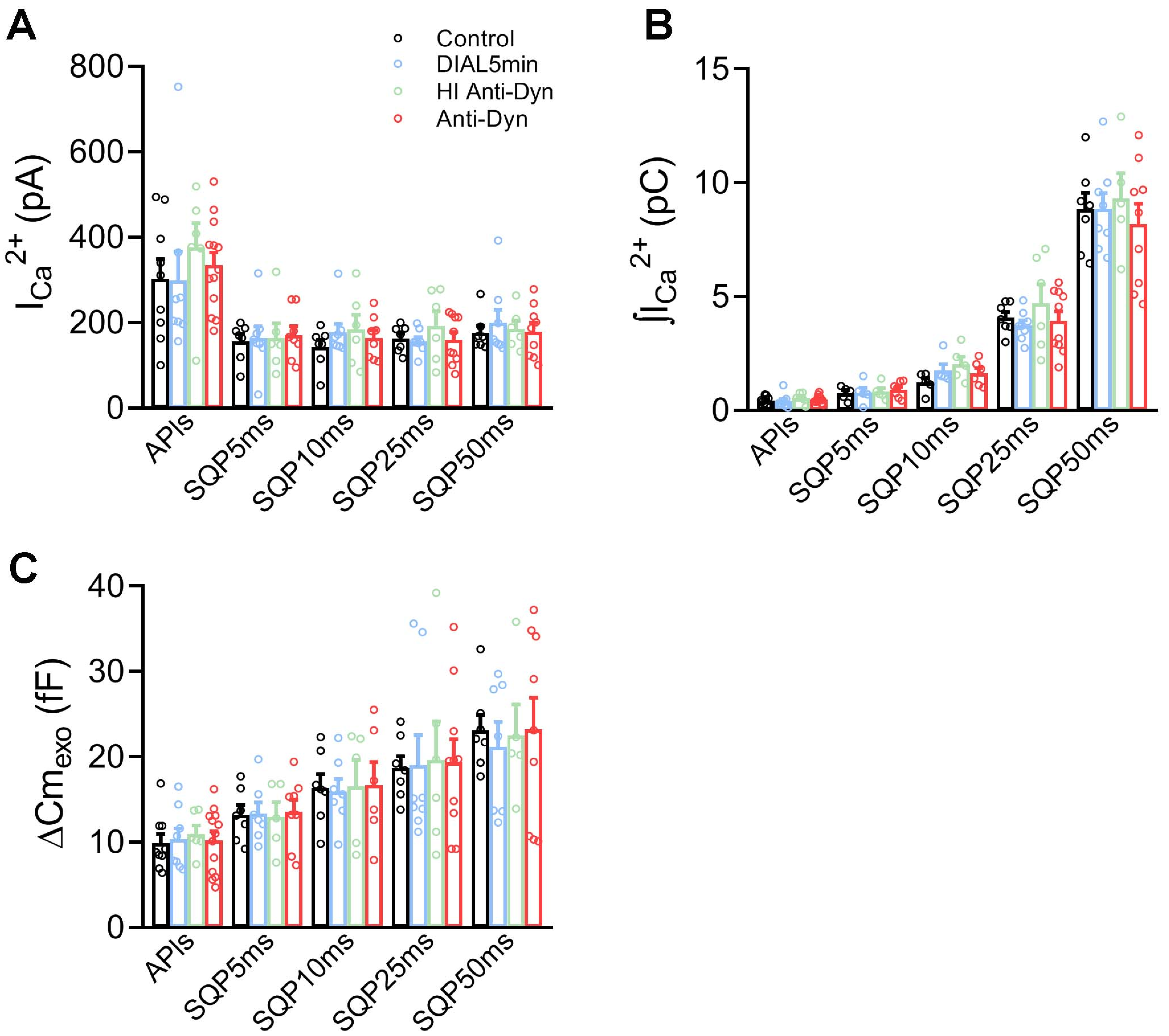


**Supplementary Figure 7:** (A), (B) and (C), The plots represent average values, standard errors and individual measurements (one measurement/cell) of $I_{Ca}2+$, $\int I_{Ca}2+$ and $\Delta Cm_{exo}$ obtained by application of APls, SQP5ms, SQP10ms, SQP25ms and SQP50ms in control conditions (standard internal solution dialyzed for 1 min before starting with measurements, Control, black), with standard internal solution dialyzed for 5 min before starting with measurements (DIAL5min, blue), with 7 nM monoclonal anti-dynamin antibody in the internal solution dialyzed for 5 min (Anti-Dyn, red), or with a heat inactivated anti-dynamin antibody in the internal solution dialyzed for 5 min before starting with measurements (HI Anti-Dyn, green), respectively. The data were analyzed by one-way ANOVA. No difference was found between conditions. The sample sizes (number of individual cells) and the number of cell culture preparations from where these cells were obtained are summarized in the Supplementary Table 1.

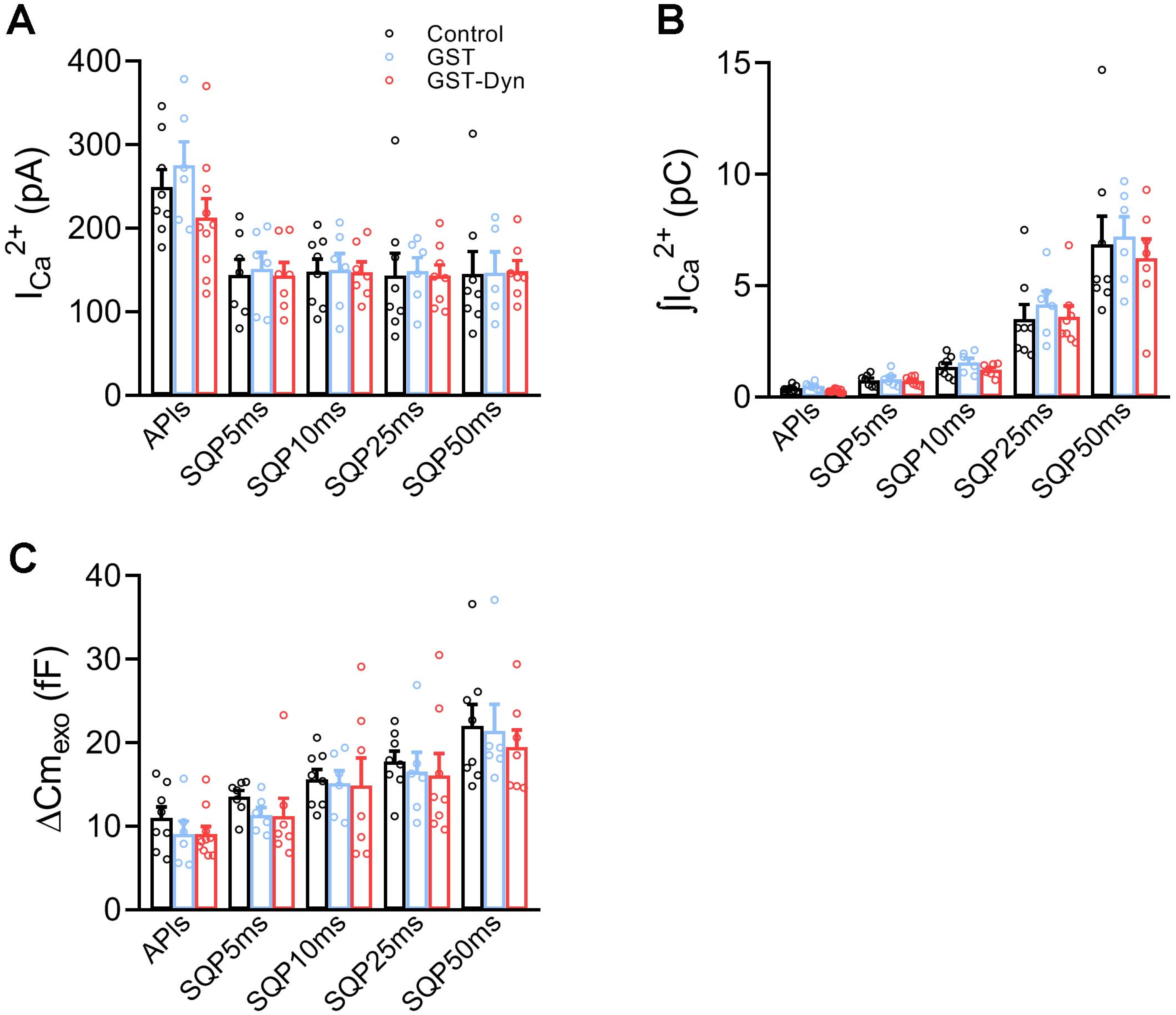


**Supplementary Figure 8:** (A), (B) and (C), The plots represent average values, standard errors and individual measurements (one measurement/cell) of $I_{Ca}$2+, $\int I_{Ca}$2+ and $\Delta Cm_{exo}$ obtained by application of APls, SQP5ms, SQP10ms, SQP25ms and SQP50ms in control conditions (standard internal solution dialyzed for 5 min before starting with measurements, Control, black), with 30 µM GST-Dyn829-842 in the internal solution dialyzed for 5 min before starting with measurements (GST-Dyn, red), or with GST in the internal solution, also dialyzed for 5 min (GST, blue), respectively. The data were analyzed by one-way ANOVA. No difference was found between conditions. The sample sizes (number of individual cells) and the number of cell culture preparations from where these cells were obtained are summarized in the Supplementary Table 1.

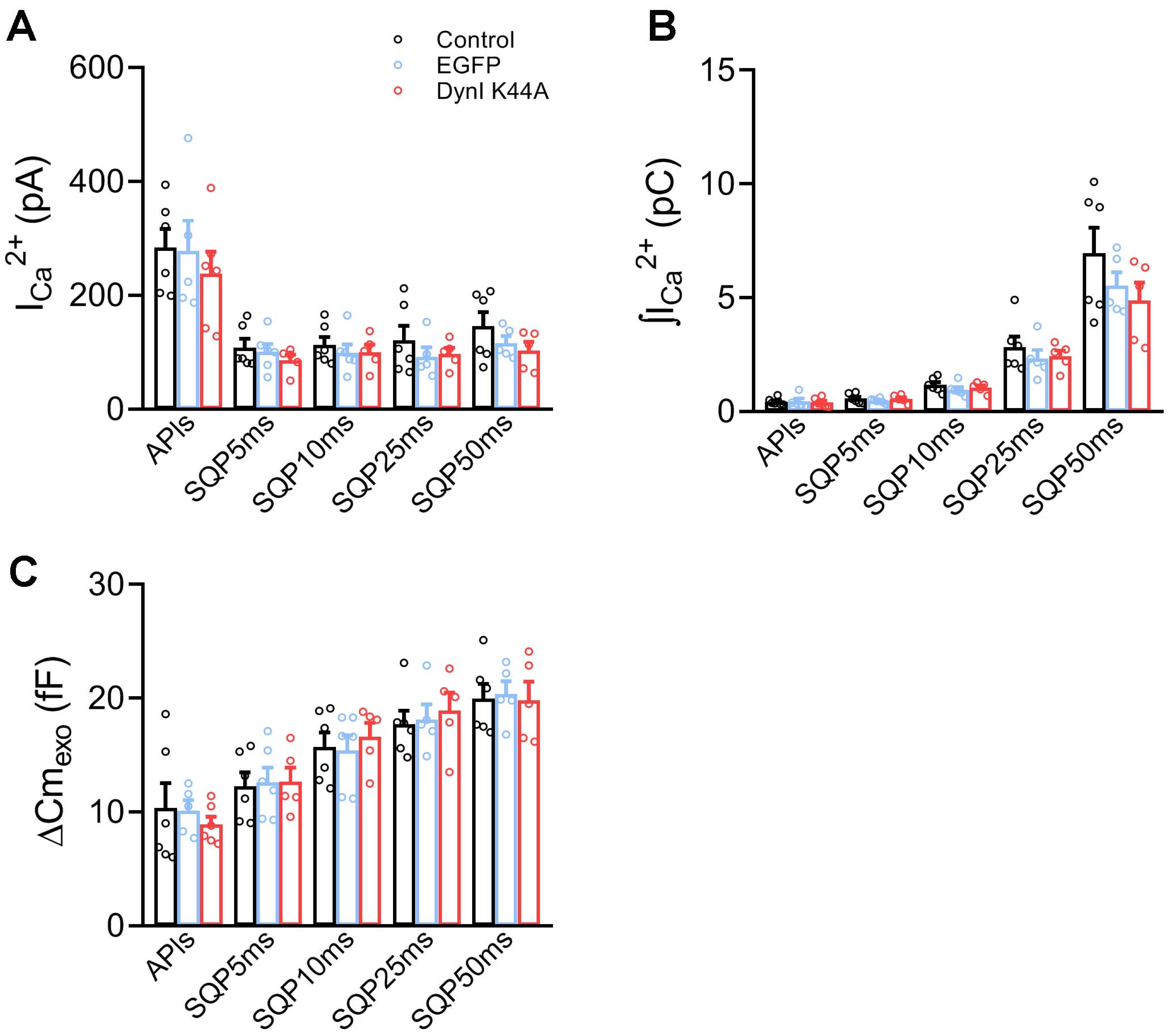


**Supplementary Figure 9:** (A), (B) and (C), The plots represent average values, standard errors and individual measurements (one measurement/cell) of $I_{Ca}2+$, $\int I_{Ca}2+$ and $\Delta Cm_{exo}$ obtained by application of APls, SQP5ms, SQP10ms, SQP25ms and SQP50ms in control conditions (Control, black), in cells expressing dominant-negative mutant dynamin I K44A-EGFP (DynI K44A, red), and cells expressing EGFP (blue), respectively. The data were analyzed by one-way ANOVA. No difference was found between conditions. The sample sizes (number of individual cells) and the number of cell culture preparations from where these cells were obtained are summarized in the Supplementary Table 1.

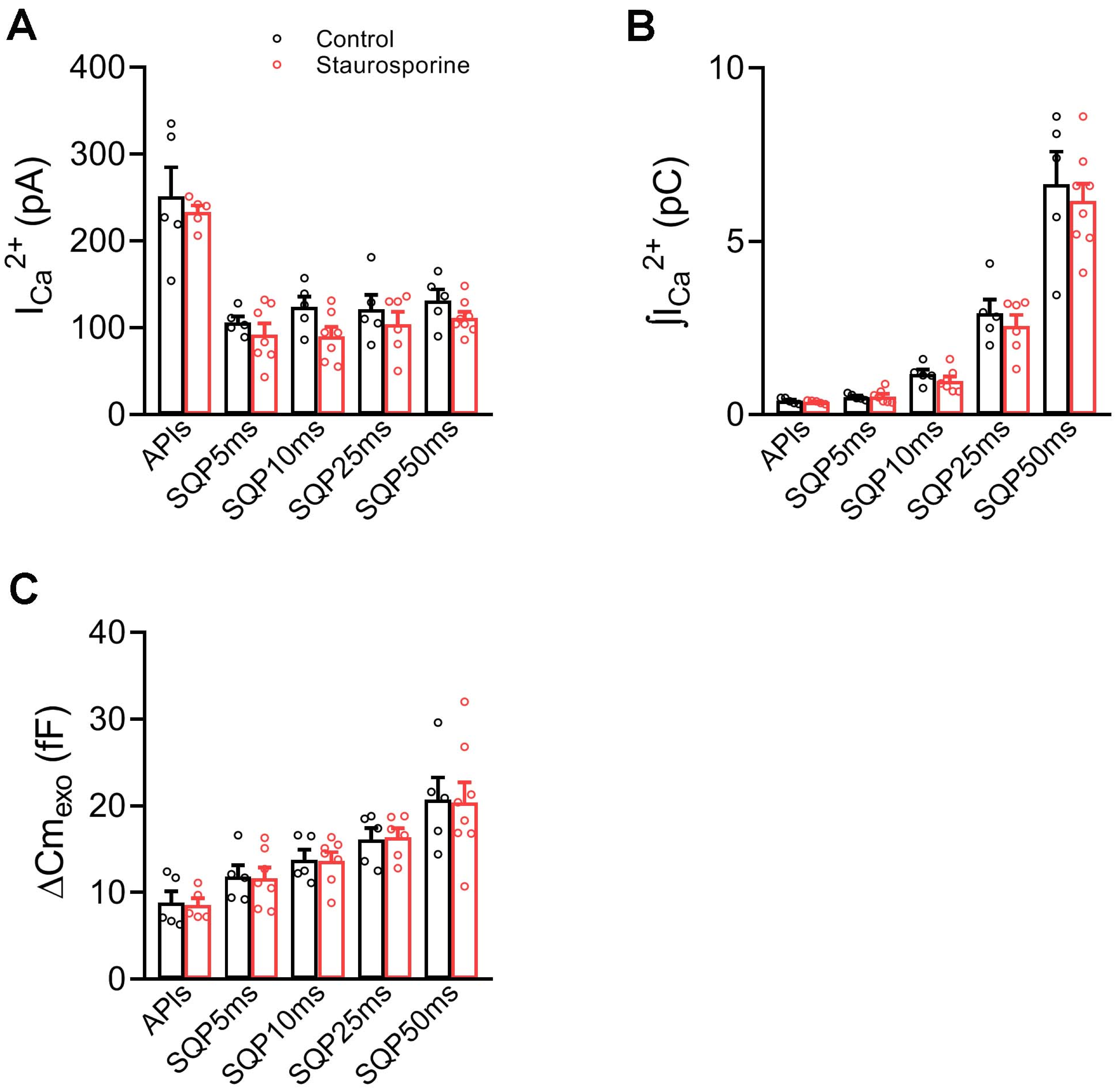


**Supplementary Figure 10:** (A), (B) and (C), The plots represent average values, standard errors and individual measurements (one measurement/cell) of $I_{Ca}2+$, $\int I_{Ca}2+$ and $\Delta Cm_{exo}$ obtained by application of APls, SQP5ms, SQP10ms, SQP25ms and SQP50ms in control conditions (Control, black) and cells treated with 100 nM staurosporine (red), respectively. The data were analyzed by Student's 't' test. No difference was found between conditions. The sample sizes (number of individual cells) and the number of cell culture preparations from where these cells were obtained are summarized in the Supplementary Table 1.

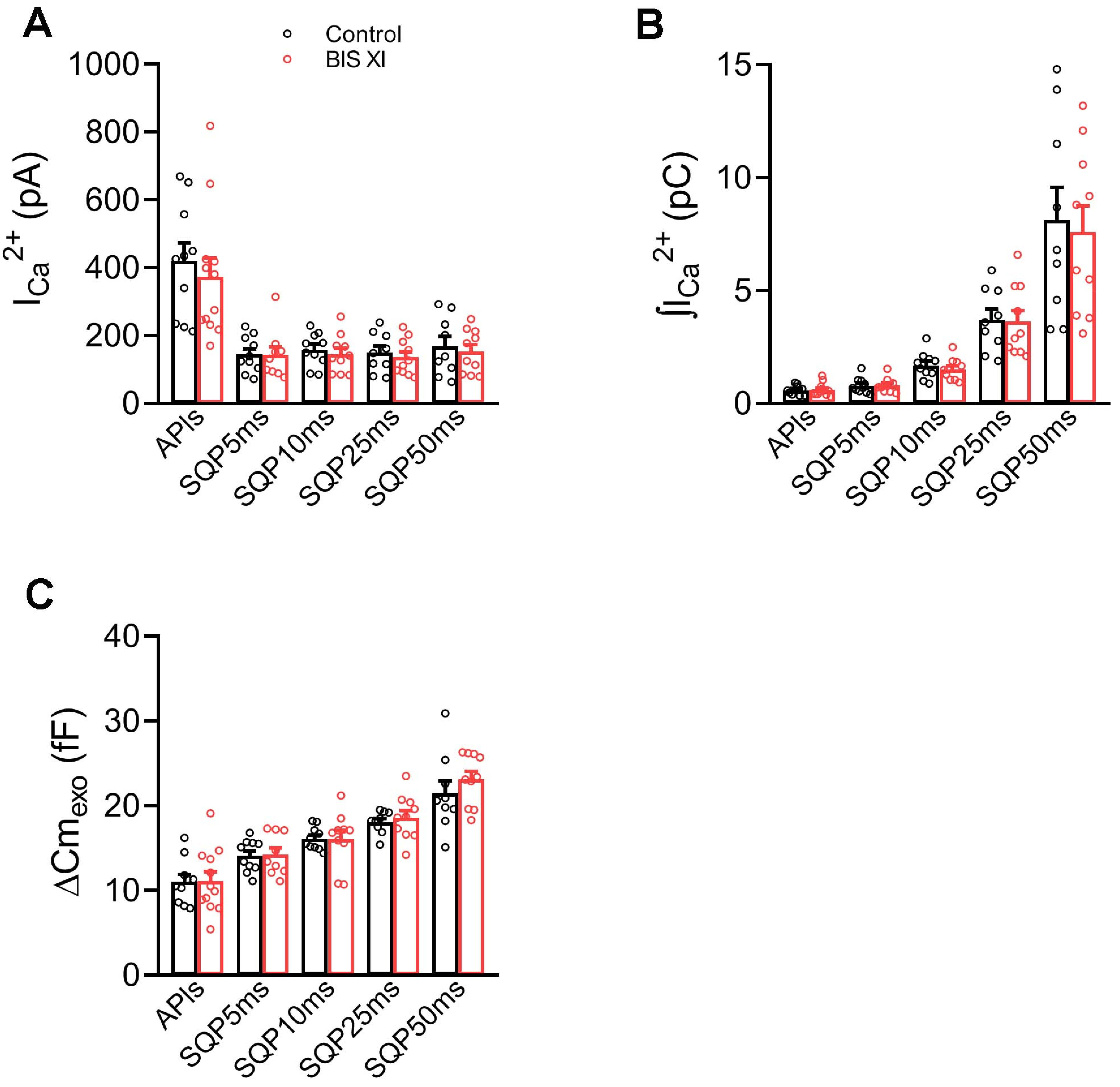


**Supplementary Figure 11:** (A), (B) and (C), The plots represent average values, standard errors and individual measurements (one measurement/cell) of $I_{Ca}2+$, $\int I_{Ca}2+$ and $\Delta Cm_{exo}$ obtained by application of APls, SQP5ms, SQP10ms, SQP25ms and SQP50ms in control conditions (Control, black) and cells treated with 3 μM bisindolylmaleimide XI (BIS XI, red), respectively. The data were analyzed by Student's 't' test. No difference was found between conditions. The sample sizes (number of individual cells) and the number of cell culture preparations from where these cells were obtained are summarized in Supplementary Table 1.

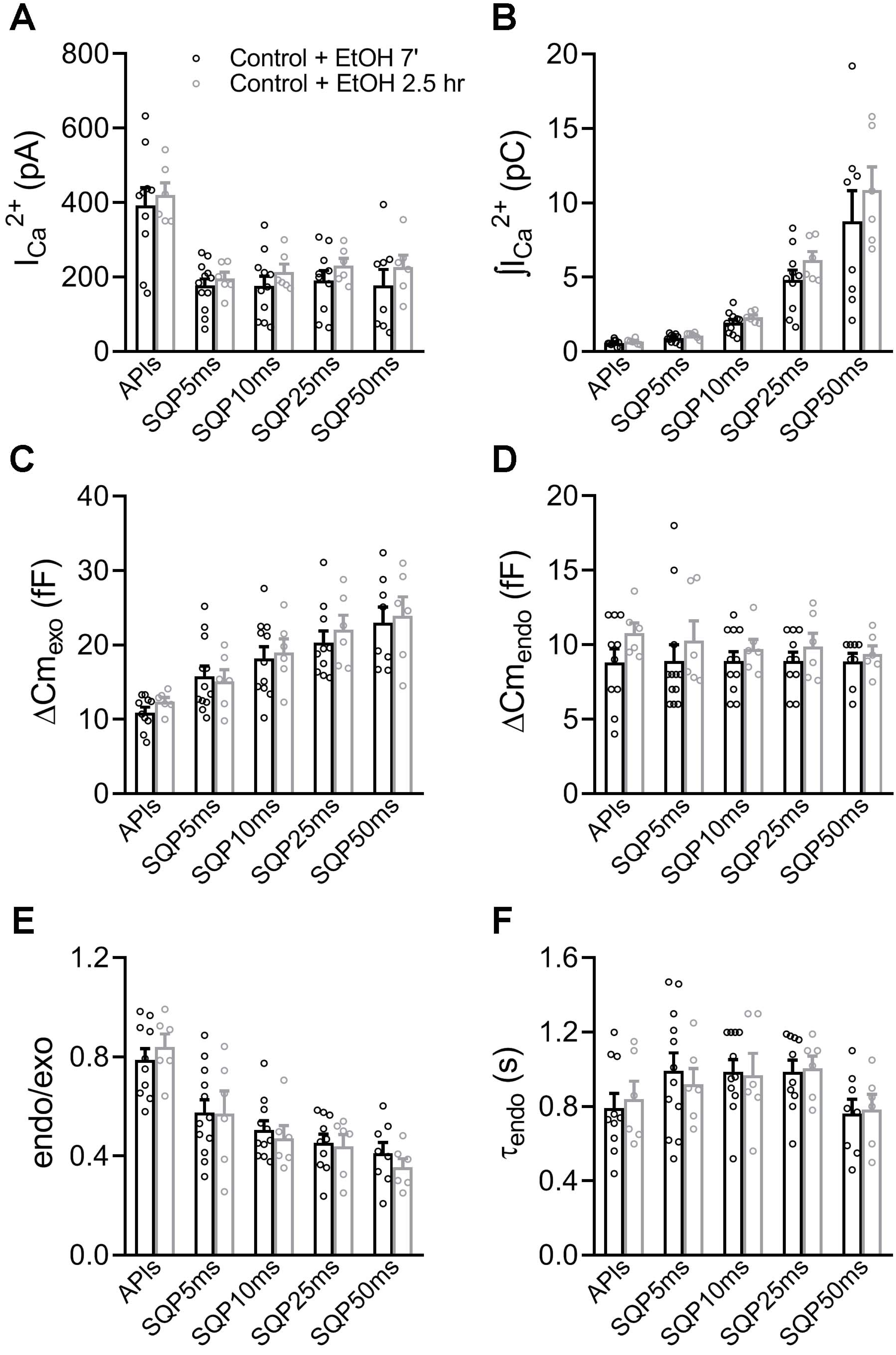


**Supplementary Figure 12:** (A), (B), (C), (D), (E) and (F). The plots represent average values, standard errors and individual measurements (one measurement/cell) of $I_{Ca}2+$, $\int I_{Ca}2+$, $\Delta Cm_{exo}$, $\Delta Cm_{endo}$, endo/exo and $\tau_{endo}$ obtained by application of APls (n=10), SQP5ms (n=12), SQP10ms (n=11), SQP25ms (n=10) and SQP50ms (n=8) in cells exposed to ethanol during 7 min at room temperature (black) and during 150 min at 37° C (gray, n=6 for all the different stimuli), respectively. The data were analyzed by Student's 't' test. No difference was found between conditions.

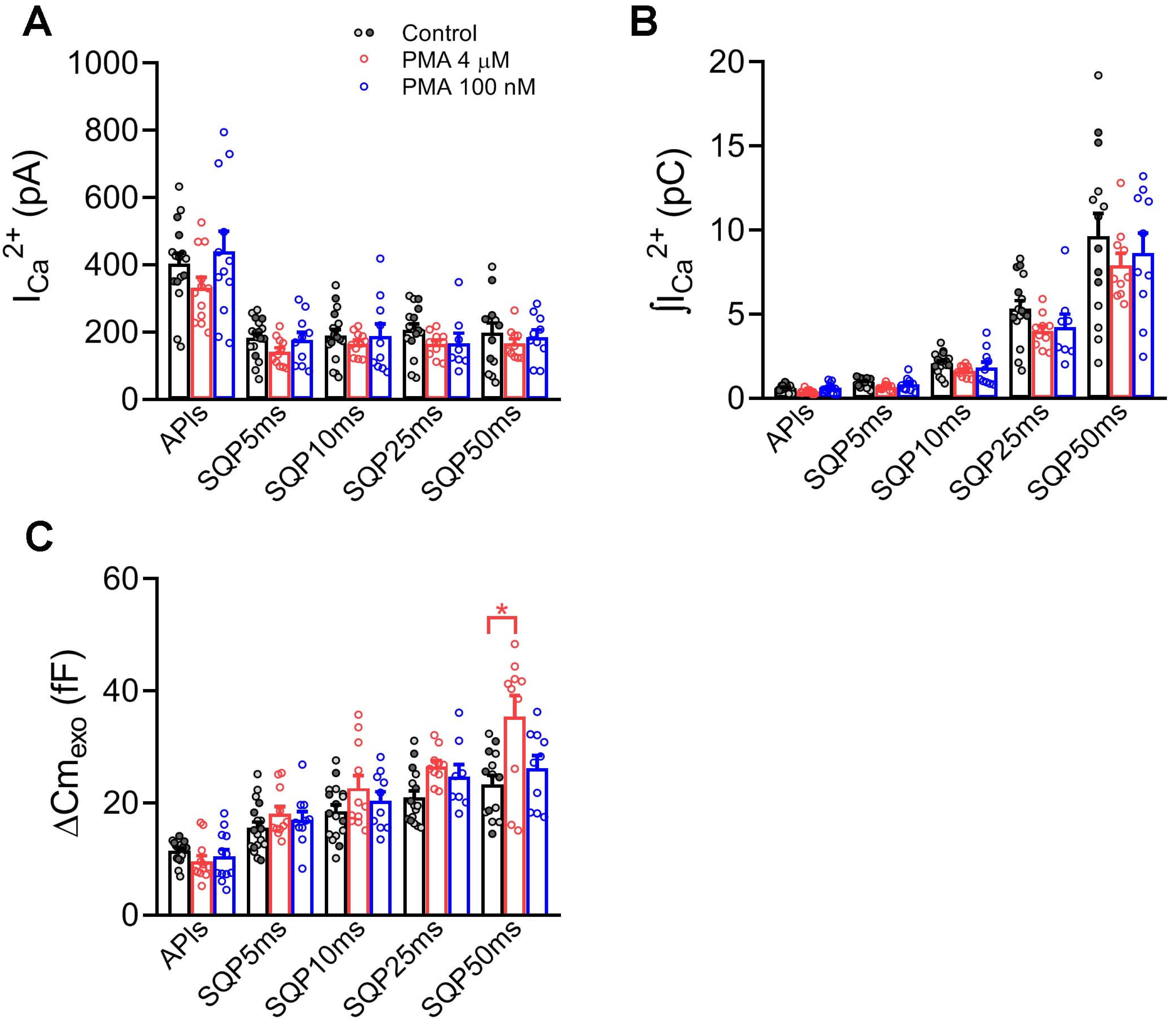


**Supplementary Figure 13:** (A), (B) and (C), The plots represent average values, standard errors and individual measurements (one measurement/cell) of $I_{Ca}2+$, $\int I_{Ca}2+$ and $\Delta Cm_{exo}$ obtained by application of APIs, SQP5ms, SQP10ms, SQP25ms and SQP50ms in Control conditions (black column, dark and light gray points), cells pretreated with 4 μM PMA (150 min, 37° C, red), and cells pretreated with 100 nM PMA (7 min, room temperature, blue), respectively. Dark and light gray points represent individual Control measurements obtained with ethanol applied during 7 min at room temperature and during 150 min at 37° C, respectively. Control column bars and SE were obtained from the grouped Control data including both conditions, which was used for statistical comparisons against both PMA treatments. The data were analyzed by one-way ANOVA followed by Bonferroni comparisons. * $p < 0.05$. The sample sizes (number of individual cells) and the number of cell culture preparations from where these cells were obtained are summarized in the Supplementary Table 1.

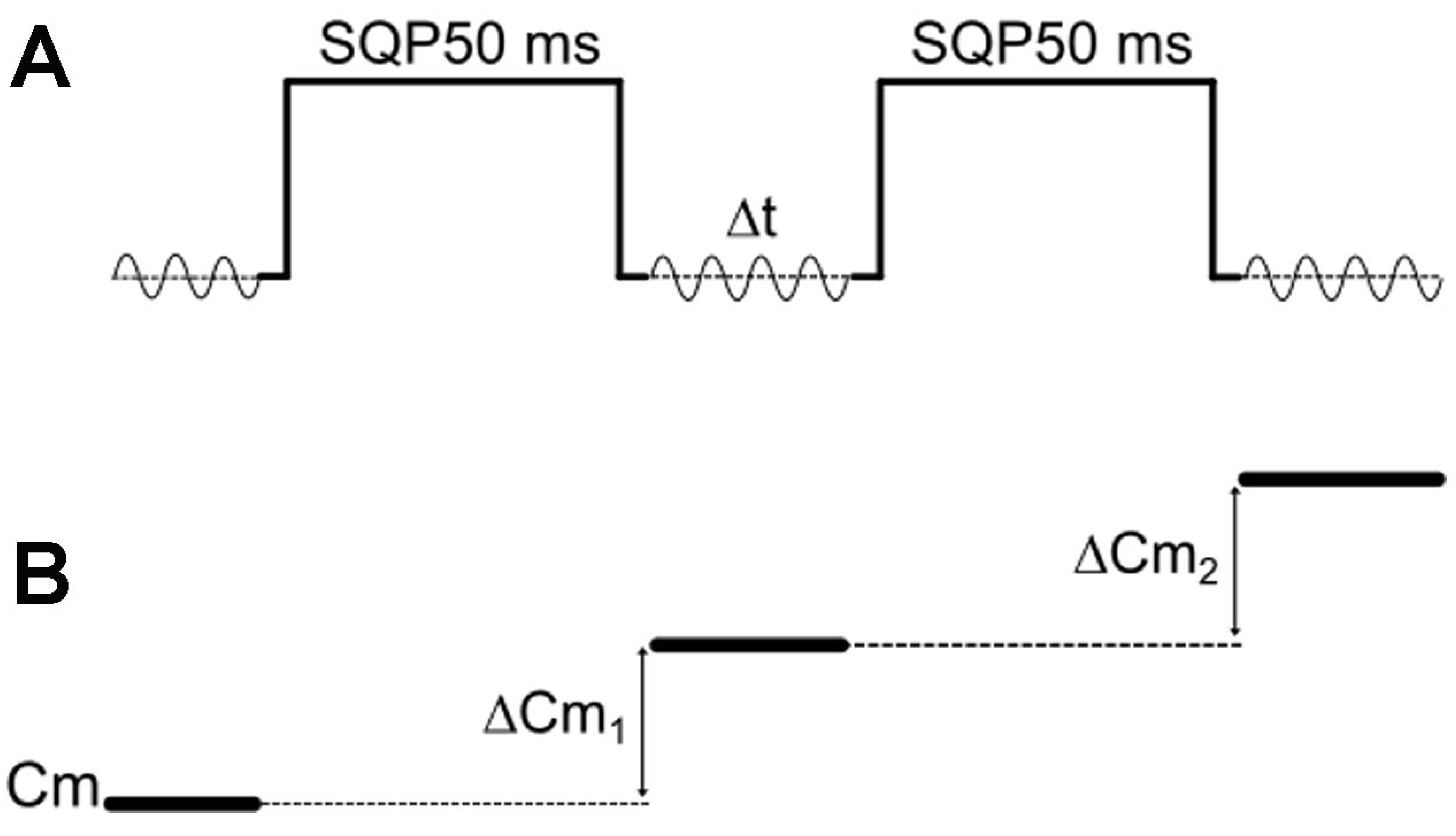


**Supplementary Figure 14:** Scheme representing the protocol of paired SQP50ms pulses (above) and the corresponding Cm record employed to evaluate IRP replenishment, as $\Delta Cm_1/\Delta Cm_2$.

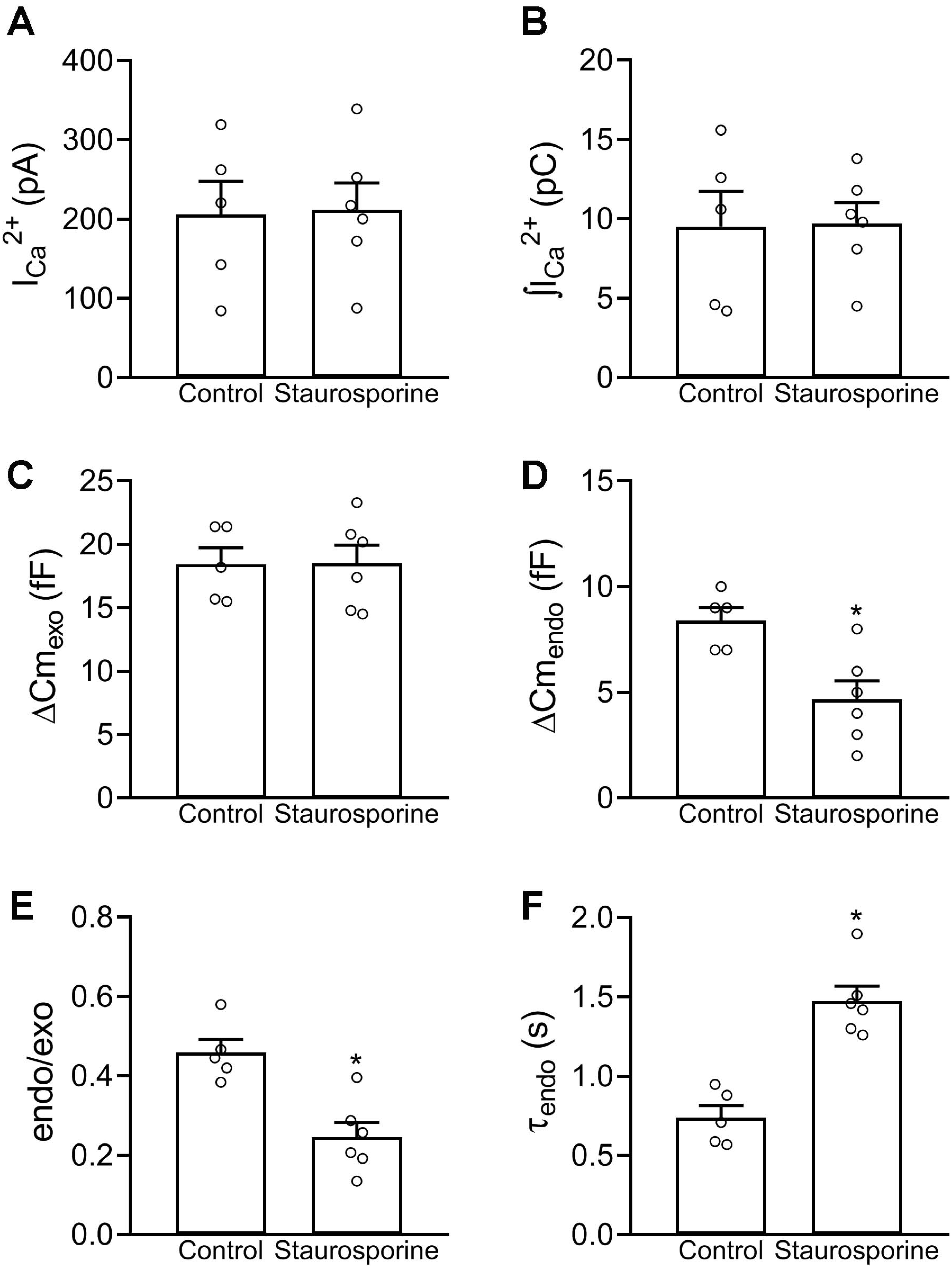


**Supplementary Figure 15:** (A), (B), (C), (D), (E) and (F). The plots represent average values, standard errors and individual measurements (one measurement/cell) of $I_{Ca}2+$, $\int I_{Ca}2+$, $\Delta Cm_{exo}$, $\Delta Cm_{endo}$, endo/exo and $\tau_{endo}$ obtained in cells stimulated with the SQP50ms paired protocol in control conditions (Control, n=5) and with 100 nM staurosporine (n=6) (see Figure 11). To standardize these variables we computed the measurements obtained on the first depolarization of the first pair of pulses applied in each cell. Additionally, to ensure completion of the endocytotic process, we only evaluated these variables in cells for which the first pair of pulses was separated with an inter-stimuli period of at least 5 s. The data were analyzed by Student's 't' test. * $p < 0.05$.

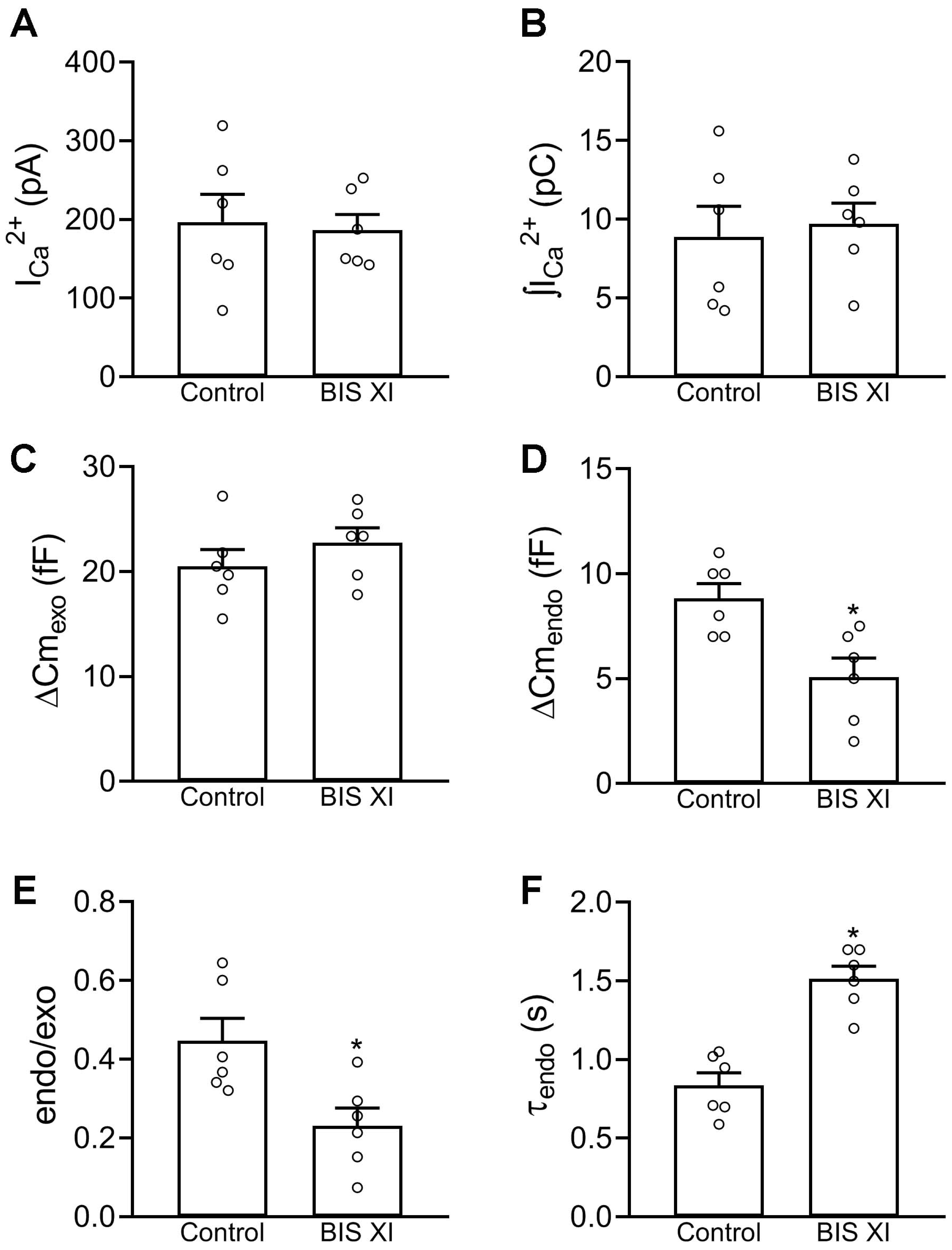


**Supplementary Figure 16:** (A), (B), (C), (D), (E) and (F), The plots represent average values, standard errors and individual measurements (one measurement/cell) of $I_{Ca}2+$, $\int I_{Ca}2+$, $\Delta Cm_{exo}$, $\Delta Cm_{endo}$, endo/exo and $\tau_{endo}$ obtained in cells stimulated with the SQP50ms paired protocol in control conditions (Control, n=6) and with 3 µM bisindolylmaleimide XI (BIS XI, n=6) (see Figure 12). To standardize these variables we computed the measurements obtained on the first depolarization of the first pair of pulses applied in each cell. Additionally, to ensure completion of the endocytic process, we only evaluated these variables in cells for which the first pair of pulses was separated with an inter-stimuli period of at least 5 s. The data were analyzed by Student's 't' test. * $p < 0.05$.

| | Nº Individual Cells (n) | | | | | Nº Cultures |
|---|---|---|---|---|---|---|
| | **APIs** | **SQP5ms** | **SQP10ms** | **SQP25ms** | **SQP50ms** | |
| **Control** | 21 | 10 | 13 | 12 | 12 | 10 |
| **$[Ca2+]_o$ 1 mM** | — | — | — | — | 7 | 7 |
| **$[Ca2+]_o$ 5 mM** | — | — | — | — | 12 | 7 |
| **$[Ca2+]_o$ 10 mM** | — | — | — | — | 10 | 7 |
| **GTPγS (Control)** | 9 | 9 | 8 | 9 | 7 | 7 |
| **GTPγS (0.3 mM)** | 13 **(9)** | 12 **(9)** | 13 **(12)** | 14 **(9)** | 9 | 7 |
| **Anti-Dyn (Control)** | 9 | 7 | 7 | 7 | 7 | 5 |
| **Anti-Dyn (DIAL 5m)** | 8 | 7 | 7 | 8 | 7 | 5 |
| **Anti-Dyn (HI Anti-Dyn)** | 6 | 5 | 5 | 6 | 5 | 5 |
| **Anti-Dyn (7 nM)** | 13 **(10)** | 8 **(7)** | 7 **(6)** | 10 | 9 | 5 |
| **GST-Dyn (Control)** | 8 | 7 | 8 | 8 | 8 | 6 |
| **GST-Dyn (GST)** | 6 | 6 | 6 | 6 | 6 | 6 |
| **GST-Dyn (30 μM)** | 10 **(5)** | 7 | 7 **(6)** | 8 | 7 | 6 |
| **Dynamin-I K44A (Control)** | 6 | 6 | 6 | 6 | 6 | 5 |
| **Dynamin-I K44A (EGFP)** | 5 | 6 | 6 | 5 | 5 | 5 |
| **Dynamin-I K44A** | 6 **(5)** | 5 | 5 | 5 | 5 | 5 |
| **Staurosporine (Control)** | 5 | 5 | 5 | 5 | 5 | 5 |
| **Staurosporine (100 nM)** | 5 | 7 | 7 | 6 | 8 **(6)** | 5 |
| **BIS XI (Control)** | 10 | 10 | 10 | 9 | 9 | 9 |
| **BIS XI (3 μM)** | 12 | 9 | 10 **(5)** | 10 **(6)** | 10 **(5)** | 9 |
| **PMA (Control 7 min & 2.5 hrs )** | 16 | 18 | 17 | 16 | 14 | 8 |
| **PMA (4 μM, 2,5 hrs)** | 12 | 11 | 11 **(6)** | 10 **(5)** | 10 **(6)** | 8 |
| **PMA (100 nM, 7 min)** | 12 | 10 | 10 | 8 | 10 | 8 |

**Table 1: Number of cells measured and number cell cultures in different experimental conditions.** In each experimental series (left, inside dark borders), all the different conditions (represented in the same colours than figures) were obtained from the same set of cultures. In some experimental conditions not all the endocytosis records were possible to be fitted to a single exponential decay equation for time constant determination. In these cases, the number between () indicates the number of cells were the time constant was estimated.
